# A resurrection study reveals limited evolution of thermal performance in response to recent climate change across the geographic range of the scarlet monkeyflower

**DOI:** 10.1101/2020.02.14.934166

**Authors:** Rachel Wooliver, Silas B. Tittes, Seema N. Sheth

## Abstract

Evolutionary rescue can prevent populations from declining under climate change, and should be more likely at high-latitude, “leading” edges of species’ ranges due to greater temperature anomalies and gene flow from warm-adapted populations. Using a resurrection study with seeds collected before and after a seven-year period of record warming, we tested for thermal adaptation in the scarlet monkeyflower *Mimulus cardinalis*. We grew ancestors and descendants from northern-edge, central, and southern-edge populations across eight temperatures. Despite recent climate anomalies, populations showed limited evolution of thermal performance curves. However, one southern population evolved a narrower thermal performance breadth by 1.25 °C, which matches the direction and magnitude of the average decrease in seasonality experienced. Consistent with the climate variability hypothesis, thermal performance breadth increased with temperature seasonality across the species’ geographic range. Inconsistent with performance trade-offs between low and high temperatures across populations, we did not detect a positive relationship between thermal optimum and mean temperature. These findings fail to support the hypothesis that evolutionary response to climate change is greatest at the leading edge, and suggest that the evolution of thermal performance is unlikely to rescue most populations from the detrimental effects of rapidly changing climate.

## Introduction

Evolution can facilitate species persistence in the face of changing climate (Hoffmann and Sgrò 2011; Carlson et al. 2014), especially when extensive habitat fragmentation prevents migration (Collingham and Huntley 2000) or plasticity is not sufficient to suit organisms to novel environments (Visser 2008). Because climate change is causing mismatches between species’ geographic ranges and thermal niches, thermal adaptation is an important driver of population responses to climate change (Geerts et al. 2015). Evolutionary rescue in the face of environmental change occurs when adaptive evolution restores positive growth rates to populations in decline, and it is most likely when the rate of environmental change is gradual and the amount of standing genetic variation for ecologically important traits is high (Carlson et al. 2014). Yet, there is severe uncertainty regarding how extreme selection events associated with changing climate (as opposed to gradual environmental change) will impact the extent to which adaptive evolution can rescue populations in decline, and whether adaptive evolution varies across species’ geographic ranges.

Capacities for thermal adaptation may vary among populations across a species’ range for at least three reasons. First, populations may experience different magnitudes of climate anomalies (departures of contemporary climate from historical averages), and thus different selective pressures on thermal tolerance. For example, temperature increases associated with climate change are often greater at higher latitudes relative to lower latitudes (IPCC 2013). Second, populations may differ in the ability to evolve earlier phenology that would enable avoidance of drought or extreme heat encountered during the growing season, consequently relaxing selection for heat tolerance (Franks et al. 2007; Sheth and Angert 2016; Socolar et al. 2017; Dickman et al. 2019). In fact, empirical studies indicate that selection for early flowering can result in correlated reductions in stress tolerance (Franks 2011; Hamann et al. 2018). Third, populations may differ in adaptive genetic variation due to connectivity with other populations. High-latitude, leading-edge populations may have ample genetic variation to evolve as they receive warm-adapted alleles from lower-latitude populations, but low-latitude, trailing-edge populations may lack genetic variation due to a scarcity of populations adapted to warmer temperatures (Davis and Shaw 2001; Hampe and Petit 2005; Hu and He 2006). Nonetheless, recent work suggests that evolutionary rescue may not occur fast enough for populations to keep up with the pace of climate change. The probability of evolutionary rescue may be especially low if amelioration of climate extremes induces reversals in trait evolution (Hamann et al. 2018) or if long generation times slow the rate of evolution (Hoffmann and Sgrò 2011). Ultimately, understanding variation in thermal niche evolution among populations could improve models that predict how species’ distributions will shift with climate change, most of which currently assume evolutionary stasis of species’ climatic niches across space and time (Angert et al. 2011; Hällfors et al. 2016; Peterson et al. 2019).

Temperature can shape species’ distributions via its effects on fitness and other performance metrics, yet we have only recently begun to understand the evolution of thermal performance across space and time (Araújo et al. 2013; Diamond 2017). A thermal performance curve (TPC) describes the performance of a genotype, population, or species across a temperature gradient (Huey and Stevenson 1979; Angilletta 2009; Fig. 1A). A TPC peaks at an intermediate temperature (thermal optimum) and is bounded by a temperature on either side where performance falls to zero (upper and lower thermal limits). The span of temperatures across which organisms achieve a designated percentage of the maximum performance is called the thermal performance breadth (hereafter referred to as breadth), and narrower breadth suggests greater thermal specialization.

**Figure 1.**
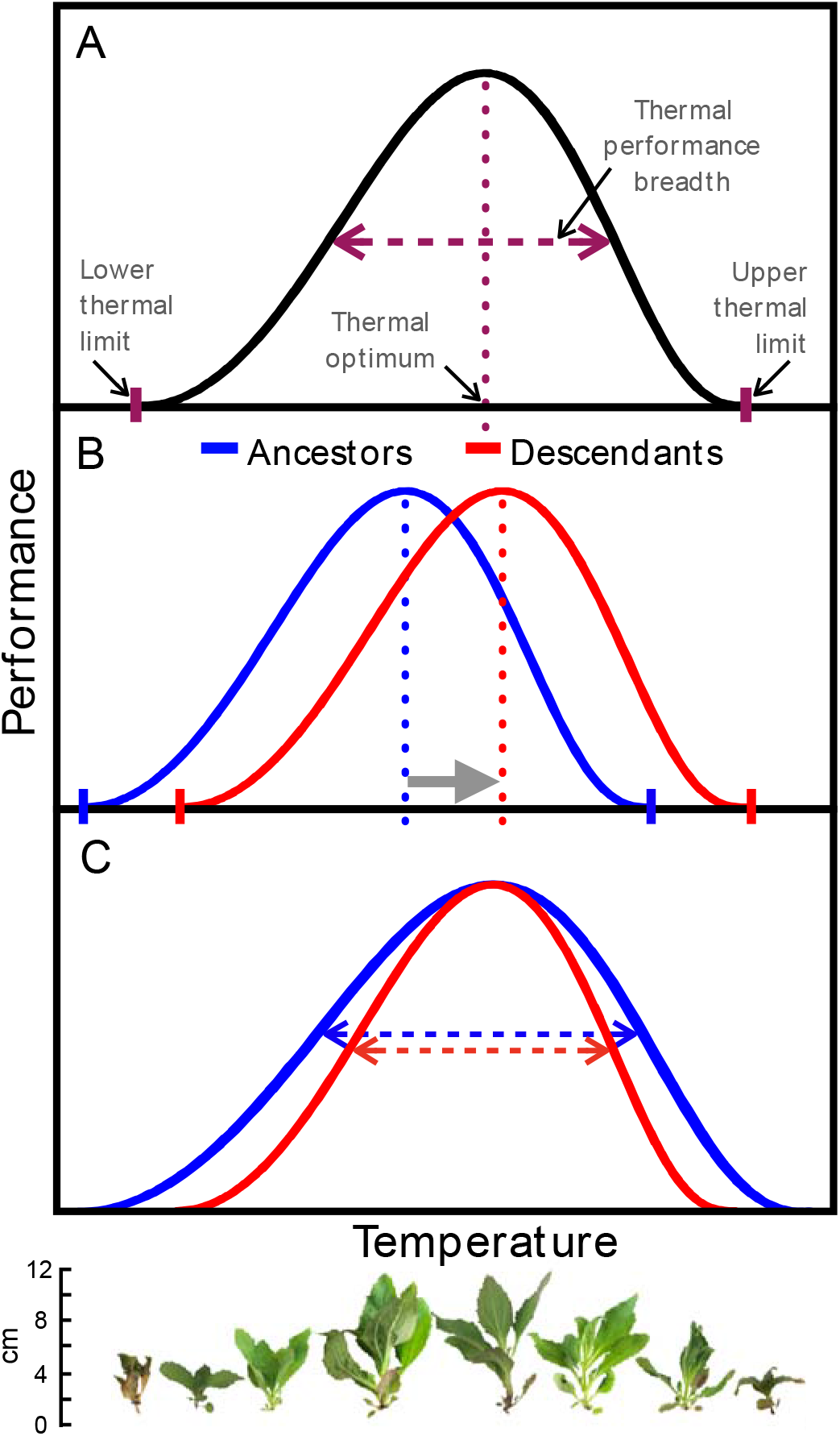
Predictions for evolution of thermal performance curves of *Mimulus cardinalis* in response to recent climate change. A) Thermal performance curves are described by parameters including the thermal optimum, thermal performance breadth, and lower and upper thermal limits. B) Thermal optima of descendants should evolve to be higher than ancestors, especially in northern and central populations where recent increases in maximum July temperatures have been most extreme, and C) breadths of descendants should evolve to be narrower than ancestors, especially in southern populations where recent decreases in temperature seasonality have been most extreme (Table 1). Images of a single S2 2010 seed family grown across the experimental temperature gradient are shown below panel C, with a scale for size.

These TPC parameters, like many other traits such as phenology or resource acquisition, can exhibit adaptive clines across spatial climatic gradients such as latitude (Lynch and Gabriel 1987; Angilletta 2009). Performance trade-offs between low and high temperatures, manifested by shifts in the TPC along the temperature axis, yield the expectation that thermal optima increase with environmental temperature (Angert et al. 2011). For example, thermal optima of populations of *Mimulus cardinalis* in western North America increased with average July temperatures and decreased with latitude, suggesting adaptive differentiation across the species’ range (Angert et al. 2011; Paul et al. 2011). If these patterns across space also apply across time, climate change-induced increases in mean temperatures should result in the evolution of increased thermal optima. The climate variability hypothesis posits that populations inhabiting regions that are climatically stable should evolve narrower climatic tolerances relative to those from climatically heterogeneous areas (Dobzhansky 1950; Janzen 1967; Stevens 1989). For ectothermic animals, thermal breadth decreases towards the equator because organisms at lower latitudes experience lower temperature variation (Sunday et al. 2010). This hypothesis has primarily been tested in temperate-tropical comparisons, but also applies to situations where climate variability changes across time, including recent climate change. That is, shifts in seasonality related to climate change could result in the evolution of altered breadth. Although there is some evidence of rapid evolution of thermal performance in response to climate change in vertebrates and invertebrates (Kingsolver et al. 2013; Higgins et al. 2014; Geerts et al. 2015), little is known about the evolution of TPCs in response to climate change in plants.

**Table 1.**
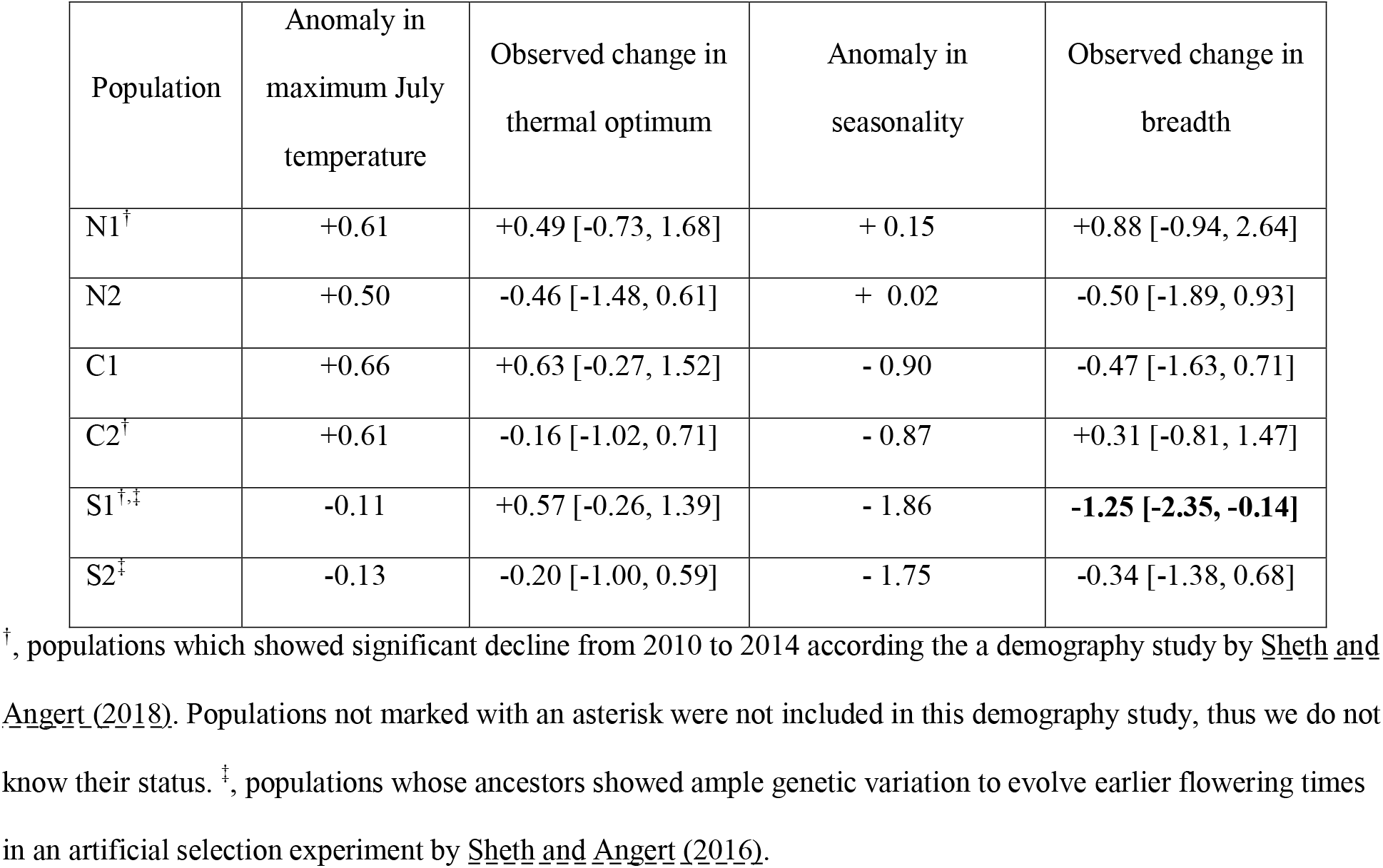
Observed evolutionary change (with 95% credible interval) in thermal optimum and breadth of six populations of *Mimulus cardinalis* (N, C, and S designate northern, central, and southern populations shown in Fig. 1A), alongside the average recent temperature anomalies (difference between historical mean and each study year from 2010 to 2017; Fig. 1B-C) for each population. Observed changes represent the mean differences in thermal optima and breadth between descendants (derived from seed collected in 2017) and ancestors (derived from seed collected in 2010) within each population. Positive values indicate that the thermal optimum or breadth was higher for descendants compared to ancestors, and negative values indicate that the thermal optimum or breadth was lower for descendants compared to ancestors. Highly plausible differences (where credible intervals do not overlap zero) are bolded. All units are in °C. Climate data were generated for population localities (Table S1) using the ClimateWNA v5.51 software package (available at http://tinyurl.com/ClimateWNA; Wang et al. 2016).

To explore how thermal performance has evolved in response to recent climate change and how the direction and magnitude of shifts vary across populations, we implemented a resurrection experiment (Franks et al. 2008) with populations across the broad geographic range of the scarlet monkeyflower, *Mimulus cardinalis* (Lowry et al. 2019). For each of two northern-edge, two central, and two southern-edge populations that collectively span broad climatic gradients in western North America (Fig. 2), we grew ancestors from 2010 alongside descendants from 2017 in growth chambers. Ancestors and descendants were respectively derived from seed collected before and after a seven-year period of record-setting drought and heat in western North America. Specifically, northern and central populations experienced the most extreme increases in temperature relative to historical conditions (Fig. 2B; Table 1). Further, temperature seasonality and annual precipitation decreased substantially in southern populations in recent years (Fig. 2C; Tables 1, S1). Recent population declines, coupled with low probabilities of survival and high probabilities of reproduction at the southern range edge, suggest that drought and warming could select for an “annualized” life history in this perennial species (Sheth and Angert 2018). Thus, decreased generation times could enhance the potential for evolutionary responses in some populations. We performed growth chamber experiments in eight temperature regimes to build TPCs for ancestors and descendants within each population. Because we held all aspects of the environment other than temperature constant, and produced seed families for both ancestors and descendants in a common environment, we can confidently attribute differences in TPCs between ancestors and descendants to genetic changes, rather than plastic developmental responses, seed storage/age effects, or maternal effects (Franks et al. 2018, 2019). We tested two hypotheses about evolutionary responses of TPCs to climate change. First, under directional warming, particularly in northern and central populations, increased thermal optima should evolve (Table 1; Fig. 1B). Second, under lower temperature seasonality, particularly in southern populations, decreased breadth should evolve (Table 1; Fig. 1C). Differences in evolutionary change in these TPC parameters among populations would suggest that thermal adaptation is dependent upon variation in climate anomalies, evolution of avoidance traits, and/or genetic variation in thermal performance. We also explored whether thermal optima and breadths are associated with geographic temperature gradients, allowing us to test for evolutionary divergence in thermal performance parameters across space.

**Figure 2.**
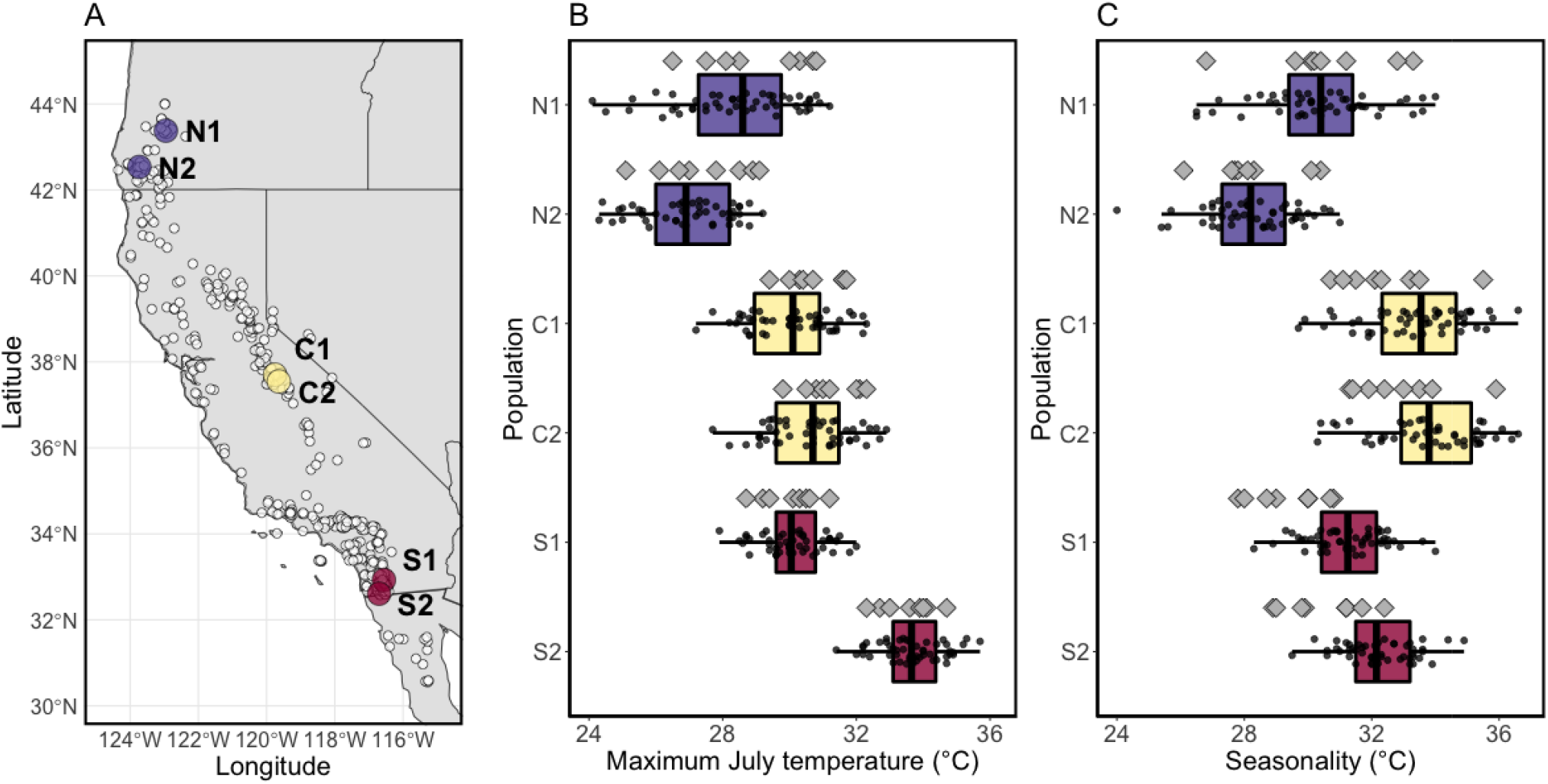
A) Map of seed collection sites of two northern-edge (purple), two central (yellow), and two southern-edge (red) populations of *Mimulus cardinalis*, superimposed on herbarium specimen locations (white circles; Angert et al. 2018). Recent (2010-2017) B) maximum July temperatures and C) temperature seasonality for each population (gray diamonds), superimposed on boxplots of historical values (1951-2000). Climate data are described in Table S1.

## Methods

### Study system, seed sampling, and refresher generation

*Mimulus cardinalis* is a perennial herb that occurs along seeps and streams from central Oregon, USA to northern Baja, Mexico (Fig. 2A). It has been the subject of numerous investigations of local adaptation, geographic range limits (Angert and Schemske 2005; Angert 2006; Paul et al. 2011; Muir and Angert 2017; Angert et al. 2018; Bayly and Angert 2019), and physiological, evolutionary, and demographic responses to climate change (Angert et al. 2011; Sheth and Angert 2016, 2018). Previous work indicates that populations in our study have been in decline, but differences in generation time and gene flow across the range may affect the potential for evolutionary rescue. Specifically, a demography study showed that during a five-year period of severe drought and warming (2010-2014), growth rates of 32 *M. cardinalis* populations decreased from the leading to trailing edges of the geographic range (Sheth and Angert 2018). Three of these populations are included in our study (N1, C2, and S1; Table 1), and each of these showed a significant population decline during the study period (Sheth and Angert 2018). Further, the demography study showed that the probability of survival from one year to the next was highest in central populations and declined towards northern and southern range edges. A majority of adults marked in 2010 in N1 and S1 populations did not survive to 2011 (Sheth and Angert 2018), and data collected beyond 2014 suggest that a few C2 plants could survive at least 6-7 years (Angert and Sheth unpubl. data). Thus, populations in the range center likely have longer generation times and lower potential for rapid evolution than those at range edges. A genetic study of the northern half of the range of *M. cardinalis* showed that northern populations have recently received a net influx of migrants from hotter environments (Paul et al. 2011), which could enhance genetic variation in thermal performance and facilitate adaptation to a warming climate.

We collected seeds from 80-100 individuals in each of the six study populations in 2010 (ancestors) and 2017 (descendants). Ancestors were collected as described in Sheth and Angert (2016), and descendants were re-collected using the same protocol. Although there is a possibility that a seedbank could have introduced individuals into the descendent populations whose parents were not exposed to the period of anomalous climate during the study period (i.e., pre-2010), previous observations have pointed towards limited seed dormancy in *M. cardinalis*. In particular, a study of mid-latitude populations (2002-2003) found that only a small fraction of seeds can remain viable in the seed bank for at least one year (Angert 2006), but a recent study of 7 populations spanning the latitudinal range (2011-2014) demonstrated that no germination occurred after the first year in the seed bank (Sheth and Angert 2018). To minimize maternal and storage/age effects, we grew seeds in the greenhouse for a ‘refresher’ generation and performed controlled crosses to produce 18 seed families within each population/cohort combination (Franks et al. 2018, 2019; Appendix S1). Most seed families had unique sires and dams (full-sibs), with the exception of some crosses that shared the same sire (half-sibs) in four population/cohort combinations with low parental sample sizes (N1 2010, N2 2010, C2 2010, and C2 2017; Table S2).

### Resurrection experiment

To determine whether *M. cardinalis* TPCs have evolved in response to recent climate change across the geographic range, we implemented a resurrection experiment in growth chambers using ancestral and descendent seed families of the six populations from the refresher generation (Appendix S1). In summary, we grew seedlings in one of eight 16h day/8h night temperature regimes (10/-5, 15/0, 20/5, 25/10, 30/15, 35/20, 40/25, or 45/30 °C) for one week. These temperature regimes encompass temperatures experienced by each of the six populations (Fig. S1) and capture full TPCs for *M. cardinalis* and close relatives (Paul et al. 2011; Sheth and Angert 2014). Previous work in *Mimulus* has documented substantial variation in growth across temperatures during the short time frame of one week (Paul et al. 2011; Sheth and Angert 2014). In each growth chamber run, we included seedlings from each of the 18 seed families within each of the 12 population/cohort combinations. Temperature regimes were replicated twice to reduce chamber effects. Each seed family was replicated four times in each temperature regime, with two replicates in each chamber run (6,912 plants total).

Prior to chamber runs, we planted seeds into 72-cell plug trays. We planted into sets of six trays, which together eventually went into each growth chamber run and contained the two replicate plants for all 216 seed families planted in a randomized design. Two to ten seeds were planted for each replicate. Seeds were germinated under a benign day/night temperature regime (20/15 °C) and a 16-hour photoperiod (6:00-22:00). Three to four weeks after planting, when most seedlings had germinated but were small enough that roots were not yet entangled, we thinned seedlings to one central-most seedling in each cell. Two weeks after thinning, when most seedlings had at least two true leaves, we put each tray set (i.e., six trays containing two replicates of all seed families) into one of four reach-in growth chambers (Percival LT-105X, Percival Scientific, Inc., Perry, Iowa, USA) that was set to one of the eight day/night temperature regimes (Appendix S1).

We measured the performance of all seedlings based on the relative change in leaf number over the course of growth chamber runs. We recorded the number of true leaves >1 mm in length on each individual immediately prior to, and one week after, being placed in the growth chamber (*leaf_in_* and *leaf_out_*, respectively). We then calculated relative growth rate (RGR) as:

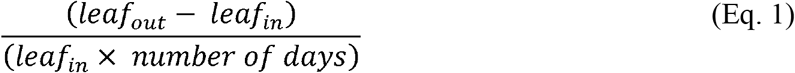

This metric does not incorporate total fitness in terms of reproductive output, and alternative performance metrics could yield different results. However, size is positively related to fruit number in natural populations of *M. cardinalis* (Sheth and Angert 2018). Because rapid growth at early life stages during which plants are smaller and more vulnerable should increase the chances of juvenile survival and thus the probability that a plant will reproduce, relative growth rate is likely correlated with total fitness. Nonetheless, we emphasize that relative growth rate is a metric of plant performance that is a component of fitness, rather than a metric of total fitness. We excluded 866 plants that died, did not germinate, or did not have true leaves by the start of chamber runs, and 13 that were accidentally damaged before final leaf number was measured. Thus, at the end of the experiment, we measured RGR for 6,033 plants (of the 6,912 initially planted). For 718 individuals that died during chamber runs (59% of plants at 10/−5 °C, 25% of plants at 45/30 °C, and <1% of plants at each of the other temperature regimes), we set RGR equal to zero. No individual had fewer leaves coming out of the chamber than going in, so all RGR values were greater than or equal to zero.

### Statistical analysis

We used RGR data to build TPCs for ancestors and descendants within each of the six *M. cardinalis* populations. To determine how thermal performance has evolved in response to recent climate change across the species’ range, we compared thermal optima and breadths of ancestors vs. descendants within each population using probabilistic comparisons (i.e., the proportion of times that the parameter for a descendent group was greater than its respective ancestral group). We used a hierarchical Bayesian model (R package *performr* v0.2; https://github.com/silastittes/performr; Tittes et al. 2019) to fit TPCs to our data. This method allowed us to simultaneously estimate responses (RGR) of our 12 biological groups (6 populations × 2 cohorts) across an environmental gradient (daytime temperature) using a derivation of Kumaraswamy’s probability density function. There are two limitations to the model in its current form. First, although there are more complex ways to model RGR as change in leaf number (Rees et al. 2010), the model is unable to handle complex response variables. Second, the model does not allow for random effects. Thus, prior to model implementation, we averaged RGR among replicates of each family in each temperature regime to avoid pseudoreplication within families and to minimize growth chamber effects (N=1,717; Table S2). We scaled RGR by the overall mean and centered daytime temperature around zero to improve model performance. We used the default model settings, except we increased iterations per chain to 10,000 and max_treedepth to 12. These settings increased convergence and reliability of posterior sampling according to 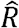 statistics of 1 and large numbers of effective samples (at least 10,000; Table S3; Gelman et al. 2014). While we did not compare our model to alternatives, we quantified the adequacy of the model’s fit to the data using a Bayesian p-value (Gelman et al. 2014). An adequate model should have a Bayesian p-value near 0.5, indicating a lack of discrepancy between the data generated by the model and the empirical data. To compute the Bayesian p-value, we compared 1) the sum of squares between the model’s expected value and the RGR values generated from each model’s posterior draws, and 2) the sum of squares when comparing posterior expectations to the empirical data. The Bayesian p-value was calculated as the proportion of posterior draws where the former sum of squares was greater than the latter. When simulating RGR values, all negative predictions were set to zero. The Bayesian p-value of our model was 0.19 (Fig. S2), indicating that the model adequately described the data generating process. Bayesian p-values for each population and cohort combination were also generally close to 0.5, though there were a few exceptions (Fig. S2). Lack of strong correlations in posterior draws between the core parameters of this model indicate that variance inflation does not influence posterior uncertainty (Fig. S3).

We derived thermal performance parameters of interest (thermal optimum and breadth) from each posterior draw of the TPC model and placed the parameters back in their original scale to aid interpretation. We calculated thermal optimum as the temperature at which RGR is maximized (Tittes et al. 2019), and breadth as the range of temperatures across which plants achieved at least 50% (B_50_) and 80% (B_80_) of maximum performance (Huey and Stevenson 1979). We calculated breadth by finding the approximate lower and upper temperature values that corresponded to 50% or 80% of the maximum height of the curve. Lacking an exact solution for the critical values, we selected them from a grid of 100 equally spaced points along the temperature axis, choosing the two points that had the minimum distance to the desired percentage of curve height. Results were qualitatively similar for B_50_ and B_80_ (Fig. S4B-C), so we report results for only B_50_ along with differences in results in cases where they influence interpretation. We performed pairwise comparisons of thermal optimum and breadth among all 12 population-by-cohort groups, focusing on comparisons between cohorts within populations. Specifically, we calculated the average and 95% credible interval of the difference in the predicted parameter estimate of ancestors vs. descendants of each population. A 95% credible interval that did not overlap zero would indicate a highly plausible difference between descendants and ancestors (i.e., we had the statistical power to detect evolutionary change). A 95% credible interval that did overlap zero would indicate that evolutionary change was not highly plausible.

To test for evolutionary divergence in thermal optima and breadth across the geographic temperature gradient, we implemented two linear models using the functions *lm* and *anova* from the *stats* package in R. We estimated means of thermal optima and breadths for each population and cohort combination from posterior draws of the TPC model. Historical maximum July temperature and historical temperature seasonality (maximum temperature of the warmest month minus minimum temperature of the coolest month) were estimated for each population and cohort combination as means from the years 1951-2000 (Table S1). The first model predicted thermal optimum as a function of maximum July temperature, cohort, and their interaction. A positive relationship between thermal optimum and maximum July temperature would confirm that there are performance trade-offs between low and high temperatures. The second model predicted breadth as a function of seasonality, cohort, and their interaction. A positive relationship between breadth and seasonality would support the climate variability hypothesis. For both models, we removed interactive and/or main effects of cohort when they were not significant at *α* < 0.05. We used R v3.6.1 for all analyses (R Core Team 2019).

## Results

### Evolution of thermal optimum

Overall, there was no support for the hypothesis that populations have evolved higher thermal optima. The thermal optimum increased by averages of about 0.5 °C from ancestors to descendants in three populations—one population from each of the northern edge (N1), central (C1), and southern edge (S1) regions of the geographic range (Figs. 3, 4A, S4A; Table 1). Thermal optimum decreased in each of the three other populations (N2, C2, and S2) by averages of less than 0.5 °C (Figs. 3, 4A, S4A; Table 1). However, because the credible intervals for all shifts in thermal optimum (both positive and negative) included 0, we inferred that evolutionary change was not highly plausible. Means and 95% credible intervals for thermal optima and other TPC parameters for all population/cohort combinations are reported in Table S4.

**Figure 3.**
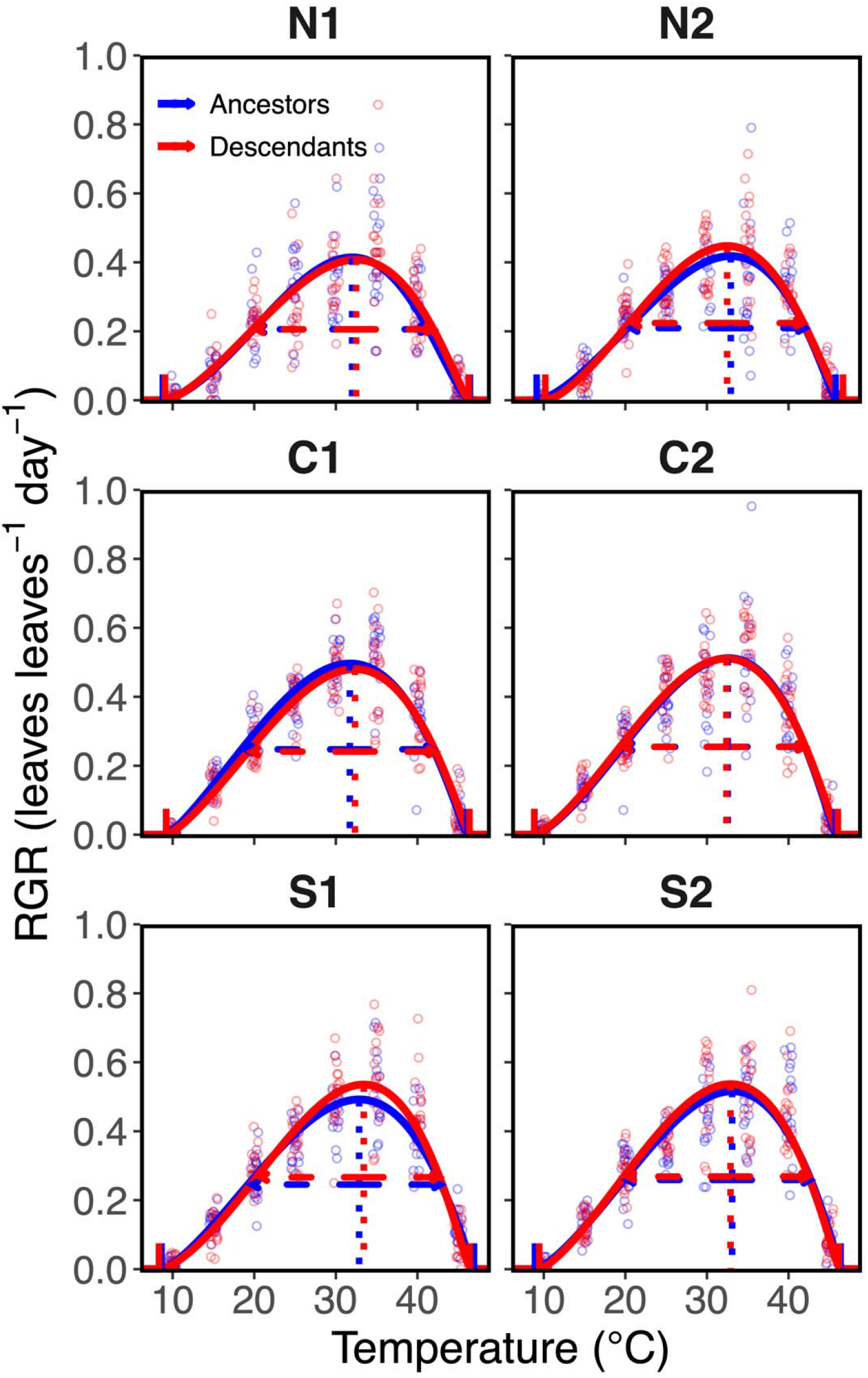
Thermal performance curves of the ancestors and descendants in each of six *Mimulus cardinalis* populations. Vertical dotted lines represent thermal optima, horizontal dashed lines represent thermal performance breadths (range of temperatures across which plants achieve 50% of maximum growth), and notches on the x-axis indicate lower and upper thermal limits. The x-axis represents daytime temperatures in growth chambers. Open points represent family means at each temperature, which are horizontally scattered around each temperature at random to aid visualization.

**Figure 4.**
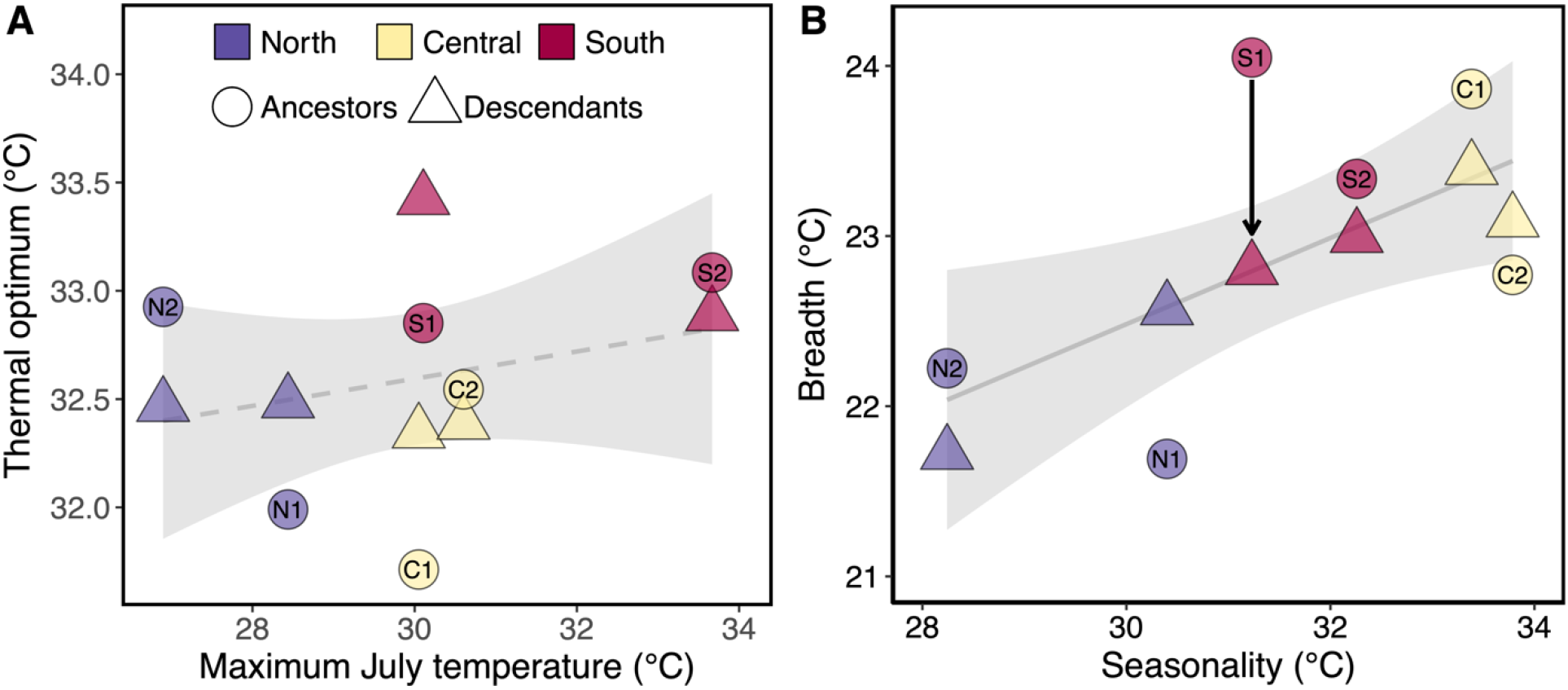
Variation in A) thermal optima across historical mean temperatures and B) breadth across historical seasonality. Thermal optima and breadth values represent mean estimates derived from a hierarchical Bayesian model, and are colored by the region where populations occur within the species’ geographic range (Fig. 2A). The arrow indicates highly plausible evolutionary change from 2010 (ancestors) to 2017 (descendants) (Table 1). Population codes are overlaid onto ancestral values. The regression in panel A was not significant at α=0.05 (dashed line), but breadth significantly increased with seasonality (solid line; see results in the main text). Shaded intervals represent 95% confidence intervals. Climate data are described in Table S1.

### Evolution of thermal performance breadth

We found mixed support for the second hypothesis that populations have evolved narrower breadths. On average, breadth increased in two populations (N1 and C2) and decreased in the other four populations (Figs. 3, 4B, S4B; Table 1). Among these, the only highly plausible evolutionary change detected was for S1, whose descendants had an average breadth that was 1.25 °C narrower than ancestors (Figs. 3, 4B, S4B; Table 1). When comparing breadth at the 80% threshold (B_80_), the direction of evolutionary change from ancestors to descendants was the same as B_50_ for each population, but that of S1 was no longer highly plausible (descendants had an average B_80_ that was 0.87 °C narrower than ancestors; Fig. S4C).

### Evolutionary divergence in thermal optimum and breadth across temperature gradients

Overall, thermal optimum did not significantly vary with maximum July temperature (*F*_1,8_=0.744, *p*=0.414) or between cohorts (*F*_1,8_=0.236, *p*=0.640). The relationship between thermal optimum and maximum July temperature did not differ between cohorts (*F*_1,8_=0.0001, *p*=0.991). After removing cohort as a main effect in the model, maximum July temperature explained no variation in thermal optimum (*b*=0.063, *R*^*2*^_*adj*_ =−0.009, *p*=0.364; Fig. 4A).

Breadth showed no significant differences between ancestors and descendants (*F*_1,8_=0.384, *p*=0.552), nor did breadth vary differently with seasonality between cohorts (*F*_1,8_=1.877, *p*=0.208). However, breadth increased with seasonality overall (*F*_1,8_=6.648, *p*=0.033). After removing cohort as a main effect in the model, seasonality explained 39% of the variation in breadth (*b*=0.253, *R*^*2*^_*adj*_ =0.386, *p*=0.018; Fig. 4B).

## Discussion

We combined a resurrection study with a hierarchical Bayesian model to test key hypotheses about how thermal performance has evolved in response to recent climate change across a plant species’ range. Since the introduction of resurrection studies over a decade ago (Franks et al. 2008), this is the first to test whether plant TPCs can rapidly evolve in response to contemporary climate change. In only seven years encompassing anomalous temperatures and record drought, we detected rapid evolution of the TPC in a southern, trailing-edge population of a perennial herb. However, we show that rapid evolution is the exception rather than the norm across multiple populations. Overall, there was no support for the hypothesis that the populations have evolved higher thermal optima, and little support for the hypothesis that the populations have evolved narrower breadths. One southern-edge population evolved a narrower breadth, indicating increased thermal specialization. There was no apparent evolutionary change in the breadth of northern-edge and central populations and the other southern-edge population. Below, we interpret these findings in light of genetic variation, natural selection, and the evolution of avoidance traits across the species’ geographic range, and we discuss their implications for evolutionary rescue.

### Genetic variation and selection

Genetic variation in thermal performance within populations and selective pressures from recent trends in climate may explain the observed evolutionary shifts in TPCs. Directional warming, estimated as the average anomaly in maximum July temperature during the study period, was greatest in the central and northern populations (Table 1; Fig. 2B). Further, anomalies in maximum July temperature steadily increased from 2010 to 2017 across populations (Fig. S5A, Appendix S2), reducing the likelihood that amelioration in climate would have induced reversals in trait shifts. Thus, upward evolutionary shifts in thermal optima (Fig. 1B) should have been greatest in the central and northern populations if they are successfully adapting to climate change. However, we did not detect significant increases in thermal optima in these populations (Table 1; Figs. 3, S4A). One interpretation of this result is that evolutionary rescue is not occurring rapidly enough for these populations to keep up with the pace of rapid climate change (Hamann et al. 2018).

There are multiple potential explanations for the lack of evolutionary responses of thermal optimum. First, climate-driven selection on thermal performance may not have been strong enough to cause significant directional change in thermal optima. Average anomalies in maximum July temperature were negative in southern populations (Table 1), so selection for higher thermal optima could have been weak. In central and northern populations, average anomalies in maximum July temperature were positive but less than 1 °C across populations. Ancestors within all four of these populations had a thermal optimum that is at least 1 °C greater than their respective historical maximum July temperatures (Fig. 4A; Tables S1, S4). Thus, ancestors were already equipped to tolerate the increased temperatures experienced throughout the study period. Three populations showed shifts in thermal optima that were opposite to their respective anomalies in maximum July temperature (N2, C2, and S1; Table 1). However, it is worth noting that C1—whose ancestors had the lowest thermal optimum of all populations (31.71 °C) and experienced the greatest average increase in maximum July temperature (+0.66 °C)—showed a similar average increase in thermal optimum (+0.63 °C; Table 1), though the credible intervals for this shift in thermal optimum overlap zero. Two other populations showed an increase in thermal optimum that matched the magnitude of increase in maximum July temperature (N1 and S2). Interestingly, although S2 did not experience maximum July temperatures during the study period that were greater than historical averages, both cohorts within S2 have a thermal optimum that is almost 1 °C less than its historical maximum July temperature average (Fig. 4A; Tables S1, S4), and thus there may still be future selection for higher thermal optimum. Given more time under elevated temperatures, N1, C1, and S2 may have the greatest potential to track further increases in mean temperature through a shift in thermal optimum.

Second, lack of gene flow from populations adapted to warmer temperatures could constrain the evolution of thermal optima across the species’ range. Northern populations have recently received an influx of migrants from central populations that occur in hotter temperatures (Paul et al. 2011). However, contemporary populations owe less than 1% of their genotypes to recent migrants (i.e., within the last two generations; Paul et al. 2011). Further, the limited gene flow from central to northern populations that may have occurred over our study period may not have introduced genotypes with higher thermal optima. This is because central populations, though they occur in hotter environments (Fig. 2A), did not have higher thermal optima than northern populations (Figs. 4A, S4A). However, on average, the southern populations in our study had higher thermal optima than central populations (Figs. 4A, S4A). One potential explanation for this pattern is limited gene flow between southern and central populations, which has preliminarily been shown by a range-wide population genetics dataset of *M. cardinalis* (J. R. Paul, T. Parchman, A. Buerkle, and A. L. Angert, unpublished manuscript). Limited gene flow between southern and central populations, paired with our finding that central populations have not evolved higher thermal optima (Table 1), suggests that gene flow from southern populations has not enhanced adaptation to warmer temperatures in central populations. Further, the southern populations in our study may not have evolved higher thermal optima because their location within the geographic range limits the opportunity to receive alleles from warmer-adapted populations. A third reason for lack of evolution of thermal optimum is that ancestors lack genetic variation in thermal optimum, a possibility that we are currently assessing. Overall, these results indicate that evolutionary rescue has not yet occurred in *M. cardinalis* populations that have declined during years of severe warming and drought. Evolution of thermal optima may not have played an important role in buffering against *M. cardinalis* population declines in response to recent climate change, but further work is needed to assess whether populations are able to evolve in the long term.

According to the climate variability hypothesis, populations that experienced the lowest temperature variation relative to historical averages should exhibit the greatest decreases in breadth (Dobzhansky 1950; Janzen 1967; Stevens 1989). When climate is stable within the lifetime of organisms, genotypes with high performance within the narrow climatic gradient are favored (Etterson 2004). In support of the climate variability hypothesis, seasonality, which represents the span of temperatures experienced during the year, was on average dramatically lower than historical conditions in the S1 population (Table 1; Fig. 2C), and this population showed plausible evolution towards thermal specialization (i.e., breadth became narrower in descendants relative to ancestors; Figs. 3, S4B). Two factors aside from decreased seasonality could have contributed to the evolution of thermal specialization in S1. First, southern populations of *M. cardinalis* have become increasingly annualized (i.e., high probabilities of flowering and low probabilities of survival from one year to the next; Sheth and Angert 2018). Annualization could enhance the rate of evolution due to shorter generation times. Second, S1 has experienced recent drought, receiving on average 111 mm less precipitation per year compared to the historical average (the strongest drought across all six populations; Table S1). Heat compounded by drought may have truncated the growing season and reduced the range of temperatures the population encounters, thus increasing selection for thermal specialization. Lower annual precipitation could also explain trends towards decreasing breadth in two other populations, S2 and C1. Average anomalies in seasonality have been small but positive in the northern populations (Table 1; Fig. 2C). Inconsistent with the climate variability hypothesis, these northern populations did not evolve broader TPCs (Table 1; Figs. 3, S4B). However, seasonality of each study year was not always greater compared to historical conditions and did not follow a unidirectional trend across the study period (Figs. 2C, S5B), so it is possible that selection for greater breadth was neither strong nor consistent across years. Further, populations whose breadth did not evolve may have lacked standing genetic variation in breadth. Overall, these results suggest that in *M. cardinalis*, breadth may be a more evolutionarily labile trait than thermal optimum, but lack of breadth evolution in response to recent climate change in a majority of populations indicates that evolutionary rescue has not occurred in populations that are declining as temperatures have become more variable.

Although there was limited evolution of thermal optimum and breadth from ancestors to descendants, we found mixed support for our hypotheses across space. Despite limited gene flow among populations (Paul et al. 2011; J. R. Paul, T. Parchman, A. Buerkle, and A. L. Angert, unpublished manuscript) and a strong gradient in mean temperature from north to south across the geographic range (Fig. 2B), we did not detect a strong pattern of adaptive divergence in thermal optima across the geographic mean temperature gradient (Fig. 4A). Interestingly, this pattern contradicts previous work showing that thermal optima of *M. cardinalis* populations increased from the northern range edge towards the range center (Angert et al. 2011; Paul et al. 2011). Further, though our results provide mixed support for the climate variability hypothesis over time (no increase in breadth in two northern populations that experienced increased seasonality, but decreased breadth in only one of four central and southern populations that experienced decreased seasonality), our data strongly support the climate variability hypothesis across the geographic temperature gradient. Contrary to the expected pattern of increased seasonality from low to high latitudes, central and southern populations experience greater seasonality and have broader TPCs than northern populations (Figs. 4B, S4B). Overall, breadth increased with historical seasonality independent of cohort (Fig. 4B), indicating that there is adaptive divergence in breadth across temperature variation, and this genetic cline is maintained with contemporary evolution.

### Evolution of avoidance vs. tolerance traits

We have shown that only one out of six *M. cardinalis* populations has responded to recent climate change through evolution of a narrower TPC (Table 1; Figs. 3, S4A-B). The populations exhibiting no TPC evolution could instead persist under climate change through the evolution of avoidance traits (Franks et al. 2007; Sheth and Angert 2016; Socolar et al. 2017; Dickman et al. 2019). At the same time, these avoidance traits, including earlier flowering, may come with the cost of lower tolerance to environmental stress. For example, when populations of *Brassica rapa* evolved earlier flowering after a multi-year drought, they concurrently evolved decreased water-use efficiency, and thus lower ability to tolerate drought conditions (Franks 2011; Hamann et al. 2018). Using the ancestral populations from our study, Sheth and Angert (2016) quantified the response to artificial selection for early and late flowering as a proxy for each population’s adaptive potential. They found that the two southern populations rapidly responded to selection on flowering time, with early- and late-flowering selection lines diverging by ~15 days. Thus, the ancestors of the southern populations had ample genetic variation to evolve earlier flowering times, potentially allowing them to avoid the extreme drought and increased temperatures that they have recently experienced (Table S1). Additionally, early-flowering selection lines, though they did not have decreased water-use efficiency, had higher specific leaf area and leaf nitrogen content, representing a partial shift toward a fast, resource-acquisitive life history (Sheth and Angert 2016). Thus, if southern populations have evolved earlier flowering in situ since 2010 (which we are currently assessing), they may have also evolved acquisitive life histories at the expense of more resource-conservative functional traits that would promote thermal tolerance. In line with this prediction, we found that the both southern populations tended to evolve narrower TPCs (though not significant in S2; Table 1; Figs. 3, S4B), meaning that the descendants did not tolerate extreme temperatures as well as their ancestors. Populations could have exhibited evolutionary shifts in other TPC parameters, including lower and upper thermal limits, critical breadth (the difference between upper and lower limit), performance maximum, and area under the performance curve. For example, if S1 evolved a more competitive growth strategy, we might predict that it has evolved a higher performance maximum. Though descendants of the southern populations on average had a higher performance maximum than their respective ancestors, these shifts were not highly plausible (Figs. 3, S4G; Table S4). In fact, with the exception of decreased breadth in S1, we did not detect highly plausible shifts in any TPC parameter for any of our study populations (Fig. S4). Thus, our results provide mixed evidence for a clear evolutionary trade-off between the ability to avoid novel environments and environmental tolerance, at least over the seven-year study period.

Though southern populations had the genetic capacity to evolve earlier flowering time, northern and central populations did not (Sheth and Angert 2016). This may hinder their abilities to avoid hotter environments via shifts in phenology. Although phenological responses to selection in a greenhouse may differ from responses to selection in the field, these previous findings suggest that evolution of TPCs may be necessary for northern and central populations to tolerate hotter, more thermally variable environments. Although we did not document the evolution of TPCs in these populations in this study, we might detect greater evolutionary responses in a resurrection experiment that uses a future set of descendent seed families. However, the greater magnitude of increase in mean temperatures and temperature seasonality, coupled with lower capacity for thermal adaptation puts northern and central populations of *M. cardinalis* at increased risk of further population decline under continued climate change.

### Caveats

A major caveat of this study is that we performed all experiments on plants at the seedling stage. Thus, although we did not detect predicted evolutionary shifts at this early life stage, such responses may still exist at later life stages in this species, for RGR or other traits such as reproductive output. We recognize that we have not quantified the full fitness curves for the ancestors and descendants of these populations, but many important TPC studies of a variety of organisms have relied on performance metrics that are partial components of total fitness (e.g., heart rate in crabs: Gaitán-Espitia et al. 2014, feeding rate in butterfly larvae: Higgins et al. 2014, growth rate in nymphalid caterpillars, sockeye salmon, and tropical tree seedlings: Brett et al. 1969; Berger et al. 2011; Cheesman and Winter 2013, locomotor activity in *Drosophila*: Latimer et al. 2011, swimming speed in tadpoles: Bartheld et al. 2017, sprinting speed in skinks and lizards: Crowley 1985; Phillips et al. 2014). Importantly, our study represents an important first step in quantifying physiological tolerances for ancestors and descendants of plants in a resurrection study, which is complementary to many other studies of *M. cardinalis* that have explored geographic range limits and responses to climate change (e.g., Angert et al. 2011; Paul et al. 2011; Sheth and Angert 2016, 2018).

Our study has several additional caveats which might limit our inferences about the evolution of TPCs in response to recent climate change. First, we maintained a constant day/night temperature regime in growth chambers, yet it is increasingly recognized that temperature variability and frequency of temperature extremes have important consequences for physiological, ecological, and evolutionary processes. Second, the cohorts in our resurrection study are derived from two sampling periods that are only seven years apart, which may not be enough time to detect evolutionary responses in a perennial plant. Based on previously collected demography data (Sheth and Angert 2018) and personal observation, we speculate that plants could have plausibly completed one to seven generations between our sampling periods. Strikingly, despite this potential drawback, we still detected rapid evolution of breadth in the S1 population, which is particularly interesting given that individuals in southern populations tend to be shorter-lived than those in northern and central populations (Sheth and Angert 2018). Third, if seeds that remained dormant in the seed bank for multiple years contributed to the descendant cohort in our study, they could slow the rate of evolutionary change (Hairston and De Stasio 1988), but the likelihood of this is low (Sheth and Angert 2018). Fourth, the hierarchical model assumes that seed families are statistically independent, however many seed families shared a parent and were thus not genetically independent (Table S2). As a result, our estimates of TPCs could be artificially more precise for those population/cohort combinations that had fewer genetically unique families (N1 2010, N2 2010, C2 2010, and C2 2017). However, we did not identify highly plausible evolutionary patterns in these populations, suggesting that greater numbers of genetically non-independent families do not inflate estimates of evolutionary response. Thus, future studies that consider additional life stages, performance metrics, temperature regimes, and sampling years are still needed to gain a comprehensive view of long-term evolutionary responses of TPCs in *M. cardinalis*.

### Conclusions and future directions

A key question in ecology and evolutionary biology is whether populations can evolve rapidly enough to keep up with the pace of climate change. Although rates of projected climate change exceeded past rates of climatic niche evolution in a macroevolutionary study of vertebrates (Quintero and Wiens 2013), we showed that in only seven years, breadth has decreased by over 1°C in a southern population of a perennial plant. This pattern of evolution may be due to a period of drought experienced in situ from 2010 to 2017, which would have truncated the growing season and reduced the range of temperatures encountered, or a trade-off in which evolution of earlier flowering comes with the cost of thermal specialization. Breadth did not significantly shift in any of the other five populations, and thermal optima did not significantly shift in any of the six populations across the species’ range, likely due to a combination insufficient time for evolutionary change, weak selection, or lack of genetic variation in thermal performance. We conclude that populations, even those that are in the same region of a geographic range (e.g., the two southern-edge populations in our study), can vary in their evolutionary responses to climate change, having important, but often overlooked, impacts on forecasts of range shifts. More importantly, our findings demonstrate that thermal performance evolution may not occur rapidly in a majority of populations, even those where it is most expected. Overall, determining the potential for population-level TPCs to evolve in response to recent climate change represents an important step forward in understanding and predicting whether evolution can rescue populations in the face of climate change.

## Acknowledgements

We are grateful to P. Beattie, J. Smith, and A. Rosvall for help with field collections in 2010, and M. Riggins, R.L. Olliff Yang, and J.E. Shay for help with field collections in 2017. We also thank A.L. Angert for providing logistical support in the field. The National Park Service, Bureau of Land Management, and California State Parks provided collection permits. We thank J. Torres, M. Wiegmann, N. Gold, C. Yurish, E. Vtipil, and E. Coughlin for assistance with pollinations, data collection, and plant care, and the North Carolina State University Phytotron & Fox Greenhouse staff for help with plant care. We thank N.C. Emery, J.W. Benning, D.N. Anstett, C.D. Muir, and two anonymous reviewers for feedback on earlier versions of this manuscript. This project was funded by the USDA National Institute of Food and Agriculture Hatch (Project ID: 1016272) and NSF DBI (Project ID: 1523866).

## Author contributions

SNS conceived of the study; RCW and SNS designed the experiment. RCW conducted the experiments and collected the data. RCW analyzed the data with the assistance of SBT. RCW and SNS wrote the first draft of the manuscript and all authors contributed to editing the final manuscript.

## References

Angert, A. L. 2006. Demography of central and marginal populations of monkeyflowers (*Mimulus cardinalis* and *M. lewisii*). Ecology 87:2014–2025.

Angert, A. L., M. Bayly, S. N. Sheth, and J. R. Paul. 2018. Testing range-limit hypotheses using range-wide habitat suitability and occupancy for the scarlet monkeyflower (*Erythranthe cardinalis*). Am. Nat. 191:E76–E89.

Angert, A. L., and D. W. Schemske. 2005. The evolution of species’ distributions: reciprocal transplants across the elevation ranges of *Mimulus cardinalis* and *M. lewisii*. Evolution 59:1671–1684.

Angert, A. L., S. N. Sheth, and J. R. Paul. 2011. Incorporating population-level variation in thermal performance into predictions of geographic range shifts. Integr. Comp. Biol. 51:733–750.

Angilletta, M. J. 2009. Thermal adaptation: a theoretical and empirical synthesis. Oxford University Press.

Araújo, M. B., F. Ferri◻Yáñez, F. Bozinovic, P. A. Marquet, F. Valladares, and S. L. Chown. 2013. Heat freezes niche evolution. Ecol. Lett. 16:1206–1219.

Bartheld, J. L., P. Artacho, and L. Bacigalupe. 2017. Thermal performance curves under daily thermal fluctuation: A study in helmeted water toad tadpoles. J. Thermal Biol. 70:80–85.

Bayly, M. J., and A. L. Angert. 2019. Niche models do not predict experimental demography but both suggest dispersal limitation across the northern range limit of the scarlet monkeyflower (*Erythranthe cardinalis*). J. Biogeogr. 46:1316–1328.

Berger, D., M. Friberg, and K. Gotthard. 2011. Divergence and ontogenetic coupling of larval behaviour and thermal reaction norms in three closely related butterflies. Proc. R. Soc. B. 278:313–320.

Brett, J. R., J. E. Shelbourn, and C. T. Shoop. 1969. Growth rate and body composition of fingerling sockeye salmon, *Oncorhynchus nerka*, in relation to temperature and ration size. J. Fish. Res. Bd. Can. 26:2363–2394.

Carlson, S. M., C. J. Cunningham, and P. A. H. Westley. 2014. Evolutionary rescue in a changing world. Trends Ecol. Evol. 29:521–530.

Cheesman, A. W., and K. Winter. 2013. Growth response and acclimation of CO2 exchange characteristics to elevated temperatures in tropical tree seedlings. J Exp Bot 64:3817–3828.

Collingham, Y. C., and B. Huntley. 2000. Impacts of habitat fragmentation and patch size upon migration rates. Ecol. Appl. 10:131–144.

Crowley, S. R. 1985. Thermal sensitivity of sprint-running in the lizard *Sceloporus undulatus*: support for a conservative view of thermal physiology. Oecologia 66:219–225.

Davis, M. B., and R. G. Shaw. 2001. Range shifts and adaptive responses to Quaternary climate change. Science 292:673–679.

Diamond, S. E. 2017. Evolutionary potential of upper thermal tolerance: biogeographic patterns and expectations under climate change. Ann. NY Acad. Sci. 1389:5–19.

Dickman, E. E., L. K. Pennington, S. J. Franks, and J. P. Sexton. 2019. Evidence for adaptive responses to historic drought across a native plant species range. Evol. Appl. 12:1569–1582.

Dobzhansky, Th. 1950. Evolution in the tropics. Am. Sci. 38:209–221.

Etterson, J. R. 2004. Evolutionary potential of *Chamaecrista fasciculata* in relation to climate change. Ii. genetic architecture of three populations reciprocally planted along an environmental gradient in the Great Plains. Evolution 58:1459–1471.

Franks, S. J. 2011. Plasticity and evolution in drought avoidance and escape in the annual plant *Brassica rapa*. New Phytol. 190:249–257.

Franks, S. J., J. C. Avise, W. E. Bradshaw, J. K. Conner, J. R. Etterson, S. J. Mazer, R. G. Shaw, and A. E. Weis. 2008. The resurrection initiative: storing ancestral genotypes to capture evolution in action. BioScience 58:870–873.

Franks, S. J., E. Hamann, and A. E. Weis. 2018. Using the resurrection approach to understand contemporary evolution in changing environments. Evol. Appl. 11:17–28.

Franks, S. J., M. R. Sekor, S. Davey, and A. E. Weis. 2019. Artificial seed aging reveals the invisible fraction: implications for evolution experiments using the resurrection approach. Evol. Ecol., doi: 10.1007/s10682-019-10007-2.

Franks, S. J., S. Sim, and A. E. Weis. 2007. Rapid evolution of flowering time by an annual plant in response to a climate fluctuation. PNAS 104:1278–1282.

Gaitán-Espitia, J. D., L. D. Bacigalupe, T. Opitz, N. A. Lagos, T. Timmermann, and M. A. Lardies. 2014. Geographic variation in thermal physiological performance of the intertidal crab *Petrolisthes violaceus* along a latitudinal gradient. J. Exp. Biol. 217:4379–4386.

Geerts, A. N., J. Vanoverbeke, B. Vanschoenwinkel, W. Van Doorslaer, H. Feuchtmayr, D. Atkinson, B. Moss, T. A. Davidson, C. D. Sayer, and L. De Meester. 2015. Rapid evolution of thermal tolerance in the water flea *Daphnia*. Nat. Clim. Change 5:665–668.

Gelman, A., J. B. Carlin, H. S. Stern, D. B. Dunson, A. Vehtari, and D. B. Rubin. 2014. Bayesian data analysis. Third Edition. Chapman and Hall, London, UK.

Hairston, N. G., and B. T. De Stasio. 1988. Rate of evolution slowed by a dormant propagule pool. Nature 336:239–242.

Hällfors, M. H., J. Liao, J. Dzurisin, R. Grundel, M. Hyvärinen, K. Towle, G. C. Wu, and J. J. Hellmann. 2016. Addressing potential local adaptation in species distribution models: implications for conservation under climate change. Ecol. Appl. 26:1154–1169.

Hamann, E., A. E. Weis, and S. J. Franks. 2018. Two decades of evolutionary changes in *Brassica rapa* in response to fluctuations in precipitation and severe drought. Evolution 72:2682–2696.

Hampe, A., and R. J. Petit. 2005. Conserving biodiversity under climate change: the rear edge matters. Ecol. Lett. 8:461–467.

Higgins, J. K., H. J. MacLean, L. B. Buckley, and J. G. Kingsolver. 2014. Geographic differences and microevolutionary changes in thermal sensitivity of butterfly larvae in response to climate. Funct. Ecol. 28:982–989.

Hoffmann, A. A., and C. M. Sgrò. 2011. Climate change and evolutionary adaptation. Nature 470:479–485.

Hu, X.-S., and F. He. 2006. Seed and pollen flow in expanding a species’ range. J. Theor. Biol. 240:662–672.

Huey, R. B., and R. D. Stevenson. 1979. Integrating thermal physiology and ecology of ectotherms: a discussion of approaches. Integr. Comp. Biol. 19:357–366.

IPCC. 2013. Climate Change 2013: The Physical Science Basis. Contribution of Working Group I to the Fifth Assessment Report of the Intergovernmental Panel on Climate Change. (eds Stocker, T.F., D. Qin, G.-K. Plattner, M. Tignor, S.K. Allen, J. Boschung, A. Nauels, Y. Xia, V. Bex and P.M. Midgley) Cambridge University Press, Cambridge, United Kingdom and New York, NY, USA.

Janzen, D. H. 1967. Why mountain passes are higher in the tropics. Am. Nat. 101:233–249.

Kingsolver, J. G., S. E. Diamond, and L. B. Buckley. 2013. Heat stress and the fitness consequences of climate change for terrestrial ectotherms. Funct. Ecol. 27:1415–1423.

Latimer, C. a. L., R. S. Wilson, and S. F. Chenoweth. 2011. Quantitative genetic variation for thermal performance curves within and among natural populations of *Drosophila serrata*. J. Evol. Biol. 24:965–975.

Lowry, D. B., J. M. Sobel, A. L. Angert, T.-L. Ashman, R. L. Baker, B. K. Blackman, Y. Brandvain, K. J. R. P. Byers, A. M. Cooley, J. M. Coughlan, M. R. Dudash, C. B. Fenster, K. G. Ferris, L. Fishman, and M. Vallejo-Marín. 2019. The case for the continued use of the genus name *Mimulus* for all monkeyflowers. TAXON.

Lynch, M. G., and W. Gabriel. 1987. Environmental tolerance. Am. Nat. 129:283–303.

Muir, C. D., and A. L. Angert. 2017. Grow with the flow: a latitudinal cline in physiology is associated with more variable precipitation in *Erythranthe cardinalis*. J. Evol. Biol. 30.

Paul, J. R., S. N. Sheth, and A. L. Angert. 2011. Quantifying the impact of gene flow on phenotype-environment mismatch: a demonstration with the scarlet monkeyflower *Mimulus cardinalis*. Am. Nat. 178:S62–S79.

Peterson, M. L., D. F. Doak, and W. F. Morris. 2019. Incorporating local adaptation into forecasts of species’ distribution and abundance under climate change. Global Change BIol. 25:775–793.

Phillips, B. L., J. Llewelyn, A. Hatcher, S. Macdonald, and C. Moritz. 2014. Do evolutionary constraints on thermal performance manifest at different organizational scales? J. Evol. Biol. 27:2687–2694.

Quintero, I., and J. J. Wiens. 2013. Rates of projected climate change dramatically exceed past rates of climatic niche evolution among vertebrate species. Ecol. Lett. 16:1095–1103.

R Core Team. 2019. R: a language and environment for statistical computing. R Foundation for Statistical Computing, Vienna.

Rees, M., C. P. Osborne, F. I. Woodward, S. P. Hulme, L. A. Turnbull, and S. H. Taylor. 2010. Partitioning the components of relative growth rate: how important is plant size variation? Am. Nat. 176:E152–E161.

Sheth, S. N., and A. L. Angert. 2016. Artificial selection reveals high genetic variation in phenology at the trailing edge of a species range. Am. Nat. 187:182–193.

Sheth, S. N., and A. L. Angert. 2018. Demographic compensation does not rescue populations at a trailing range edge. PNAS 115:2413–2418.

Sheth, S. N., and A. L. Angert. 2014. The evolution of environmental tolerance and range size: a comparison of geographically restricted and widespread *Mimulus*. Evolution 68:2917–2931.

Socolar, J. B., P. N. Ebanchin, S. R. Beissinger, and M. W. Tingley. 2017. Shifts in time and space interact as climate warms. PNAS 114:12976–12981.

Stevens, G. C. 1989. The latitudinal gradient in geographical range: how so many species coexist in the tropics. Am. Nat. 133:240–256.

Sunday, J. M., A. E. Bates, and N. K. Dulvy. 2010. Global analysis of thermal tolerance and latitude in ectotherms. Proc. R. Soc. B. 278:1823–1830.

Thompson, R. M., J. Beardall, J. Beringer, M. Grace, and P. Sardina. 2013. Means and extremes: building variability into community-level climate change experiments. Ecol. Lett. 16:799–806.

Tittes, S. B., J. F. Walker, L. Torres-Martínez, and N. C. Emery. 2019. Grow where you thrive, or where only you can survive? An analysis of performance curve evolution in a clade with diverse habitat affinities. Am. Nat. 193:530–544.

Visser, M. E. 2008. Keeping up with a warming world; assessing the rate of adaptation to climate change. Proc. R. Soc. B. 275:649–659.

Wang, T., A. Hamann, D. Spittlehouse, and C. Carroll. 2016. Locally downscaled and spatially customizable climate data for historical and future periods for North America. PLoS ONE 11:e0156720.

