## Supporting Information for "A resurrection study reveals limited evolution of thermal performance in response to recent climate change across the geographic range of the scarlet monkeyflower"

**Table S1**. GPS coordinates, elevation, and climatic data for the six *Mimulus cardinalis* populations included in the resurrection study. All historical climate averages were calculated based on annual climate estimates from 1951 to 2000. Temperature seasonality was calculated as the maximum temperature of the warmest month minus minimum temperature of the coolest month. Climate data were generated for localities of each of the six populations using the ClimateWNA v5.51 software package (available at http://tinyurl.com/ClimateNWA; Wang et al*.* 2016, cited in the main text).

|  |  |  |  | **Historical average / average anomaly** | | | |
| --- | --- | --- | --- | --- | --- | --- | --- |
| **Population** | **Latitude (°N)** | **Longitude (°W)** | **Elevation (m)** | **Mean annual temperature (°C)** | **Maximum July temperature (°C)** | **Annual precipitation (mm)** | **Temperature seasonality (°C)** |
| N1 | 43.37876 | -122.95207 | 295 | 11.00 / 0.80 | 28.44 / 0.61 | 1559 / 40.22 | 30.40 / 0.15 |
| N2 | 42.53529 | -123.73016 | 914 | 10.17 / 0.74 | 26.90 / 0.50 | 1730 / 38.82 | 28.24 / 0.02 |
| C1 | 37.70377 | -119.75363 | 1316 | 11.75 / 1.16 | 30.05 / 0.66 | 951 / -16.09 | 33.38 / -0.90 |
| C2 | 37.54576 | -119.64152 | 1228 | 11.96 / 1.30 | 30.60 /0.61 | 1057 / -20.2 | 33.79 / -0.87 |
| S1 | 32.92788 | -116.56019 | 1252 | 12.81 / 0.94 | 30.11 / -0.11 | 726 / -110.91 | 31.23 / -1.86 |
| S2 | 32.60831 | -116.70098 | 252 | 16.57 / 0.93 | 33.67 / -0.13 | 366 / -53.62 | 32.26 / -1.75 |

**Table S2**. Number of full-sib and half-sib seed families for each population and cohort used in the thermal performance experiment, and sample sizes of each population and cohort for family-averaged data used in the Bayesian thermal performance model. Full-sib families have unique dams and sires, while half-sib families have unique dams but share a sire with one or more other seed families. Number of unique sires for each population and cohort is also shown.

| **Population and year** | **Number of full sib families** | **Number of half sib families** | **Number of unique sires** | **Sample size** |
| --- | --- | --- | --- | --- |
| N1 2010 | 1 | 17 | 7 | 137 |
| N1 2017 | 18 | 0 | 18 | 144 |
| N2 2010 | 10 | 8 | 14 | 144 |
| N2 2017 | 18 | 0 | 18 | 144 |
| C1 2010 | 18 | 0 | 18 | 144 |
| C1 2017 | 18 | 0 | 18 | 144 |
| C2 2010 | 1 | 17 | 8 | 140 |
| C2 2017 | 2 | 16 | 9 | 144 |
| S1 2010 | 18 | 0 | 18 | 144 |
| S1 2017 | 18 | 0 | 18 | 144 |
| S2 2010 | 18 | 0 | 18 | 144 |
| S2 2017 | 18 | 0 | 18 | 144 |

**Table S3**. Stan model fit summary. Columns represent the summary statistics, posterior percentiles, effective number of samples (“n_eff ”), and $\hat{R}$ (“R-hat”) statistic for the marginal posterior of each parameter (rows; parameters are described in the legend of Fig. S3). Indices in square brackets correspond to the 12 population-by-cohort combinations, ordered by population within cohort (N1 2010 [1], N2 2010 [2], C1 2010 [3], C2 2010 [4], S1 2010 [5], S2 2010 [6], N1 2017 [7], N2 2017 [8], C1 2017 [9], C2 2017 [10], S1 2017 [11], and S2 2017 [12]). The large number of effective samples (out of 20,000 total) indicate efficient sampling of the posterior and near-independent samples (i.e., low autocorrelation). An $\hat{R}$ value near or at 1 suggests that the model is appropriately sampling the posterior distribution.

|  | mean | se_mean | sd | 2.50% | 25% | 50% | 75% | 97.50% | n_eff | Rhat |
| --- | --- | --- | --- | --- | --- | --- | --- | --- | --- | --- |
| shape1[1] | 2.56 | 0 | 0.16 | 2.26 | 2.45 | 2.55 | 2.66 | 2.87 | 10795.18 | 1 |
| shape1[2] | 2.45 | 0 | 0.11 | 2.25 | 2.38 | 2.45 | 2.52 | 2.67 | 15370.28 | 1 |
| shape1[3] | 2.22 | 0 | 0.06 | 2.11 | 2.18 | 2.22 | 2.26 | 2.36 | 14421.51 | 1 |
| shape1[4] | 2.39 | 0 | 0.08 | 2.26 | 2.34 | 2.39 | 2.44 | 2.57 | 14765.09 | 1 |
| shape1[5] | 2.33 | 0 | 0.07 | 2.21 | 2.28 | 2.32 | 2.37 | 2.47 | 15431.95 | 1 |
| shape1[6] | 2.37 | 0 | 0.07 | 2.23 | 2.32 | 2.36 | 2.41 | 2.52 | 17659.98 | 1 |
| shape1[7] | 2.43 | 0 | 0.13 | 2.2 | 2.34 | 2.42 | 2.51 | 2.69 | 13263.41 | 1 |
| shape1[8] | 2.45 | 0 | 0.15 | 2.16 | 2.34 | 2.45 | 2.55 | 2.75 | 11704.66 | 1 |
| shape1[9] | 2.31 | 0 | 0.08 | 2.16 | 2.25 | 2.31 | 2.36 | 2.49 | 13671.08 | 1 |
| shape1[10] | 2.36 | 0 | 0.08 | 2.23 | 2.31 | 2.36 | 2.41 | 2.54 | 13186.58 | 1 |
| shape1[11] | 2.5 | 0 | 0.09 | 2.34 | 2.44 | 2.49 | 2.56 | 2.69 | 15237.34 | 1 |
| shape1[12] | 2.35 | 0 | 0.07 | 2.22 | 2.3 | 2.34 | 2.39 | 2.49 | 15901.61 | 1 |
| shape2[1] | 2.5 | 0 | 0.27 | 2.07 | 2.31 | 2.48 | 2.67 | 3.09 | 15455.87 | 1 |
| shape2[2] | 2.12 | 0 | 0.11 | 2 | 2.04 | 2.09 | 2.16 | 2.39 | 21020.54 | 1 |
| shape2[3] | 2.1 | 0 | 0.09 | 2 | 2.03 | 2.08 | 2.14 | 2.32 | 16947.29 | 1 |
| shape2[4] | 2.1 | 0 | 0.09 | 2 | 2.04 | 2.08 | 2.15 | 2.33 | 17361.25 | 1 |
| shape2[5] | 2.09 | 0 | 0.08 | 2 | 2.03 | 2.07 | 2.13 | 2.31 | 16377.92 | 1 |
| shape2[6] | 2.05 | 0 | 0.05 | 2 | 2.02 | 2.04 | 2.07 | 2.19 | 25704.36 | 1 |
| shape2[7] | 2.23 | 0 | 0.18 | 2.01 | 2.09 | 2.19 | 2.32 | 2.67 | 17314.68 | 1 |
| shape2[8] | 2.4 | 0 | 0.23 | 2.04 | 2.22 | 2.37 | 2.54 | 2.92 | 15270.25 | 1 |
| shape2[9] | 2.13 | 0 | 0.11 | 2 | 2.05 | 2.1 | 2.19 | 2.42 | 16779.51 | 1 |
| shape2[10] | 2.13 | 0 | 0.11 | 2 | 2.05 | 2.1 | 2.18 | 2.4 | 14921.05 | 1 |
| shape2[11] | 2.1 | 0 | 0.1 | 2 | 2.03 | 2.08 | 2.15 | 2.35 | 20412.04 | 1 |
| shape2[12] | 2.07 | 0 | 0.07 | 2 | 2.02 | 2.05 | 2.1 | 2.25 | 22132.50 | 1 |
| stretch[1] | 0.87 | 0 | 0.04 | 0.79 | 0.84 | 0.87 | 0.89 | 0.95 | 20083.92 | 1 |
| **Table S3**. *(continued)* |  |  |  |  |  |  |  |  |  |  |
|  | mean | se_mean | sd | 2.50% | 25% | 50% | 75% | 97.50% | n_eff | Rhat |
| stretch[2] | 0.92 | 0 | 0.03 | 0.86 | 0.9 | 0.92 | 0.94 | 0.99 | 22327.58 | 1 |
| stretch[3] | 1.15 | 0 | 0.03 | 1.09 | 1.13 | 1.15 | 1.17 | 1.21 | 26326.41 | 1 |
| stretch[4] | 1.14 | 0 | 0.03 | 1.07 | 1.11 | 1.14 | 1.16 | 1.2 | 25963.88 | 1 |
| stretch[5] | 1.11 | 0 | 0.03 | 1.05 | 1.09 | 1.11 | 1.13 | 1.17 | 25683.62 | 1 |
| stretch[6] | 1.16 | 0 | 0.04 | 1.09 | 1.14 | 1.16 | 1.19 | 1.24 | 27099.05 | 1 |
| stretch[7] | 0.9 | 0 | 0.04 | 0.83 | 0.88 | 0.9 | 0.93 | 0.98 | 22064.70 | 1 |
| stretch[8] | 0.97 | 0 | 0.04 | 0.89 | 0.94 | 0.97 | 1 | 1.05 | 14565.52 | 1 |
| stretch[9] | 1.09 | 0 | 0.03 | 1.03 | 1.07 | 1.09 | 1.12 | 1.16 | 22705.23 | 1 |
| stretch[10] | 1.14 | 0 | 0.03 | 1.08 | 1.12 | 1.14 | 1.16 | 1.21 | 22261.14 | 1 |
| stretch[11] | 1.16 | 0 | 0.04 | 1.09 | 1.13 | 1.16 | 1.18 | 1.23 | 24576.06 | 1 |
| stretch[12] | 1.21 | 0 | 0.03 | 1.15 | 1.19 | 1.21 | 1.23 | 1.28 | 25005.13 | 1 |
| nu[1] | 10.83 | 0 | 0.73 | 9.42 | 10.34 | 10.83 | 11.32 | 12.28 | 33755.87 | 1 |
| nu[2] | 13.03 | 0 | 0.83 | 11.44 | 12.46 | 13.02 | 13.59 | 14.69 | 33313.40 | 1 |
| nu[3] | 17.02 | 0.01 | 1.08 | 14.95 | 16.29 | 17.01 | 17.73 | 19.15 | 33273.13 | 1 |
| nu[4] | 16.56 | 0.01 | 1.05 | 14.52 | 15.84 | 16.55 | 17.26 | 18.66 | 33544.43 | 1 |
| nu[5] | 18.29 | 0.01 | 1.15 | 16.09 | 17.5 | 18.28 | 19.06 | 20.57 | 37194.49 | 1 |
| nu[6] | 14.77 | 0.01 | 0.95 | 12.93 | 14.12 | 14.77 | 15.41 | 16.66 | 34417.57 | 1 |
| nu[7] | 11.42 | 0 | 0.76 | 9.94 | 10.9 | 11.41 | 11.93 | 12.93 | 33702.23 | 1 |
| nu[8] | 14.02 | 0 | 0.9 | 12.27 | 13.4 | 14.01 | 14.63 | 15.81 | 35770.04 | 1 |
| nu[9] | 14.73 | 0.01 | 0.94 | 12.93 | 14.09 | 14.73 | 15.35 | 16.6 | 32988.54 | 1 |
| nu[10] | 16.1 | 0.01 | 1.02 | 14.15 | 15.4 | 16.09 | 16.78 | 18.13 | 33475.28 | 1 |
| nu[11] | 14.8 | 0 | 0.96 | 12.96 | 14.15 | 14.8 | 15.43 | 16.68 | 37738.64 | 1 |
| nu[12] | 16.91 | 0.01 | 1.08 | 14.82 | 16.17 | 16.91 | 17.63 | 19.05 | 35295.14 | 1 |
| x_min[1] | -18.73 | 0.01 | 0.86 | -20.32 | -19.28 | -18.77 | -18.26 | -16.75 | 11889.98 | 1 |
| x_min[2] | -18.39 | 0.01 | 0.69 | -19.72 | -18.83 | -18.41 | -17.98 | -16.92 | 15223.78 | 1 |
| x_min[3] | -18.33 | 0 | 0.3 | -18.99 | -18.5 | -18.3 | -18.12 | -17.82 | 15145.37 | 1 |
| x_min[4] | -18.67 | 0 | 0.42 | -19.58 | -18.92 | -18.64 | -18.38 | -17.94 | 15138.35 | 1 |
| x_min[5] | -18.89 | 0 | 0.37 | -19.7 | -19.12 | -18.86 | -18.62 | -18.25 | 17119.68 | 1 |
| x_min[6] | -18.6 | 0 | 0.45 | -19.56 | -18.88 | -18.57 | -18.29 | -17.81 | 17388.60 | 1 |
| x_min[7] | -18.46 | 0.01 | 0.71 | -19.9 | -18.92 | -18.45 | -18.01 | -17.01 | 15463.93 | 1 |
| x_min[8] | -17.33 | 0.01 | 0.94 | -19.07 | -18.02 | -17.35 | -16.66 | -15.48 | 12635.75 | 1 |
| x_min[9] | -18.3 | 0 | 0.45 | -19.24 | -18.58 | -18.29 | -18.02 | -17.42 | 14403.58 | 1 |
| x_min[10] | -18.69 | 0 | 0.41 | -19.6 | -18.94 | -18.65 | -18.4 | -17.98 | 14560.49 | 1 |
| x_min[11] | -19.2 | 0 | 0.52 | -20.3 | -19.53 | -19.17 | -18.84 | -18.28 | 16050.30 | 1 |
| x_min[12] | -18.08 | 0 | 0.4 | -18.88 | -18.33 | -18.08 | -17.84 | -17.23 | 15477.39 | 1 |
| x_max[1] | 18.91 | 0 | 0.53 | 18.09 | 18.52 | 18.85 | 19.23 | 20.11 | 18603.95 | 1 |
| x_max[2] | 18.25 | 0 | 0.22 | 17.94 | 18.1 | 18.21 | 18.36 | 18.8 | 21046.51 | 1 |
| x_max[3] | 18.58 | 0 | 0.22 | 18.26 | 18.42 | 18.53 | 18.69 | 19.12 | 19988.24 | 1 |
| x_max[4] | 18.26 | 0 | 0.18 | 18.01 | 18.14 | 18.23 | 18.35 | 18.72 | 19758.01 | 1 |
| x_max[5] | 19.41 | 0 | 0.31 | 18.96 | 19.19 | 19.35 | 19.56 | 20.16 | 21758.03 | 1 |
| **Table S3**. *(continued)* |  |  |  |  |  |  |  |  |  |  |
|  | mean | se_mean | sd | 2.50% | 25% | 50% | 75% | 97.50% | n_eff | Rhat |
| x_max[6] | 18.86 | 0 | 0.21 | 18.53 | 18.71 | 18.83 | 18.98 | 19.35 | 23739.33 | 1 |
| x_max[7] | 18.76 | 0 | 0.41 | 18.2 | 18.46 | 18.69 | 18.98 | 19.78 | 19735.26 | 1 |
| x_max[8] | 19.15 | 0 | 0.51 | 18.35 | 18.76 | 19.09 | 19.46 | 20.31 | 17638.18 | 1 |
| x_max[9] | 18.86 | 0 | 0.32 | 18.42 | 18.64 | 18.8 | 19.02 | 19.64 | 20086.21 | 1 |
| x_max[10] | 18.52 | 0 | 0.25 | 18.18 | 18.34 | 18.47 | 18.64 | 19.13 | 16792.81 | 1 |
| x_max[11] | 18.89 | 0 | 0.28 | 18.49 | 18.7 | 18.84 | 19.03 | 19.59 | 23390.33 | 1 |
| x_max[12] | 18.65 | 0 | 0.19 | 18.36 | 18.51 | 18.62 | 18.75 | 19.13 | 23899.67 | 1 |

**Table S4**. Means and 95% credible intervals of thermal performance curve parameters for each population and cohort in the resurrection experiment. All estimates except for area have been back-transformed to their original scales, as RGR was scaled by the overall mean and temperature was centered around zero prior to analysis. All units are in °C except performance maximum (leaves leaves^-1^ day^-1^).

|  | Thermal performance parameter | | | | | | | |
| --- | --- | --- | --- | --- | --- | --- | --- | --- |
| Population and year | Thermal optimum | B50 | B80 | Critical breadth | Lower thermal limit | Upper thermal limit | Performance maximum | Area |
| N1 2010 | 31.99 [31.15, 32.90] | 21.69 [20.33, 23.11] | 13.15 [12.15, 14.23] | 37.64 [35.34, 39.72] | 8.79 [7.20, 10.77] | 46.43 [45.61, 47.63] | 0.41 [0.37, 0.45] | 32.57 [30.19, 35.05] |
| N1 2017 | 32.48 [31.63, 33.33] | 22.57 [21.39, 23.65] | 13.79 [12.87, 14.63] | 37.23 [35.59, 39.10] | 9.06 [7.62, 10.51] | 46.29 [45.72, 47.30] | 0.41 [0.38, 0.45] | 33.61 [31.31, 35.97] |
| N2 2010 | 32.93 [32.21, 33.61] | 22.22 [21.26, 23.10] | 13.59 [12.83, 14.26] | 36.65 [35.13, 38.12] | 9.13 [7.80, 10.60] | 45.78 [45.46, 46.33] | 0.42 [0.39, 0.45] | 33.68 [31.66, 35.78] |
| N2 2017 | 32.47 [31.71, 33.26] | 21.72 [20.65, 22.77] | 13.24 [12.39, 14.10] | 36.48 [34.20, 38.82] | 10.19 [8.45, 12.04] | 46.67 [45.88, 47.83] | 0.45 [0.42, 0.48] | 35.34 [33.35, 37.37] |
| C1 2010 | 31.71 [31.11, 32.28] | 23.86 [23.05, 24.58] | 14.75 [14.08, 15.33] | 36.90 [36.27, 37.80] | 9.20 [8.54, 9.70] | 46.10 [45.78, 46.64] | 0.50 [0.47, 0.52] | 42.37 [40.45, 44.35] |
| C1 2017 | 32.34 [31.65, 33.00] | 23.39 [22.48, 24.24] | 14.39 [13.65, 15.04] | 37.17 [36.15, 38.47] | 9.22 [8.28, 10.10] | 46.38 [45.94, 47.16] | 0.48 [0.45, 0.51] | 40.61 [38.51, 42.79] |
| C2 2010 | 32.54 [31.93, 33.12] | 22.77 [21.94, 23.49] | 13.96 [13.28, 14.53] | 36.93 [36.12, 38.01] | 8.86 [7.95, 9.58] | 45.78 [45.54, 46.24] | 0.51 [0.48, 0.54] | 41.93 [39.87, 44.05] |
| C2 2017 | 32.39 [31.74, 32.99] | 23.09 [22.21, 23.85] | 14.17 [13.46, 14.79] | 37.21 [36.38, 38.40] | 8.84 [7.93, 9.54] | 46.04 [45.71, 46.65] | 0.51 [0.48, 0.54] | 42.46 [40.41, 44.56] |
| S1 2010 | 32.85 [32.27, 33.39] | 24.05 [23.29, 24.80] | 14.79 [14.18, 15.37] | 38.30 [37.50, 39.43] | 8.63 [7.82, 9.27] | 46.93 [46.48, 47.68] | 0.49 [0.46, 0.52] | 42.53 [40.74, 44.40] |
| S1 2017 | 33.42 [32.79, 34.02] | 22.80 [21.99, 23.61] | 13.92 [13.27, 14.55] | 38.09 [37.07, 39.44] | 8.32 [7.22, 9.24] | 46.42 [46.01, 47.11] | 0.53 [0.50, 0.57] | 44.13 [41.82, 46.52] |
| S2 2010 | 33.08 [32.49, 33.64] | 23.34 [22.57, 24.10] | 14.33 [13.73, 14.94] | 37.46 [36.63, 38.48] | 8.92 [7.97, 9.71] | 46.38 [46.05, 46.87] | 0.52 [0.48, 0.55] | 43.57 [41.24, 46.01] |
| S2 2017 | 32.89 [32.31, 33.42] | 22.99 [22.28, 23.66] | 14.13 [13.55, 14.68] | 36.73 [35.84, 37.65] | 9.44 [8.64, 10.30] | 46.17 [45.88, 46.65] | 0.54 [0.51, 0.57] | 44.51 [42.40, 46.71] |

*Thermal optimum: temperature at which plant performance is maximized; B50: span of temperatures across which plants achieve at least 50% of maximum performance; B80: span of temperatures across which plants achieve at least 80% of maximum performance. Critical breadth: span of temperatures between lower and upper thermal limits; Lower thermal limit: minimum temperature at which performance falls to 0; Upper thermal limit: maximum temperature at which performance falls to 0; Performance maximum: maximum performance achieved across temperatures; Area: area under the thermal performance curve.*

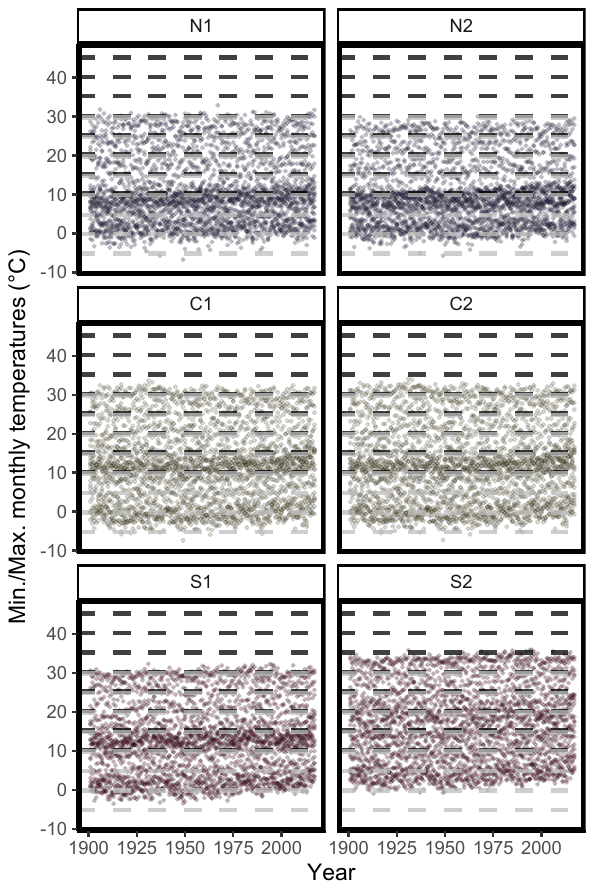

**Figure S1**. Minimum and maximum monthly mean temperatures for six populations of *Mimulus cardinalis* (purple: northern edge, yellow: central, red: southern edge). Values are shown across time (1900-2017), superimposed on temperature regimes implemented in the growth chamber study (black dashed lines: daytime temperatures, gray dashed lines: nighttime temperatures). Climate data were generated for population localities (Table S1) using the ClimateWNA v5.51 software package (available at <http://tinyurl.com/ClimateWNA>; Wang et al. 2016, cited in the main text).

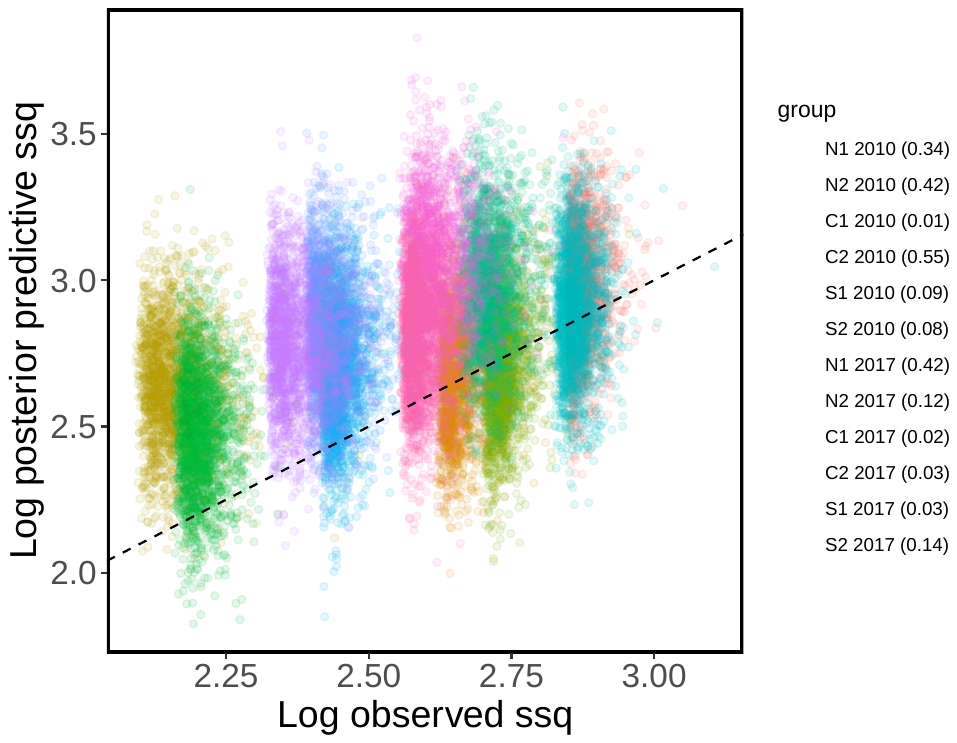

**Figure S2**. Empirical and posterior predictive sum of squares (ssq) for each population and cohort combination, with a one-to-one line. Values in parentheses represent each group’s Bayesian p-value. The overall Bayesian p-value was 0.19.

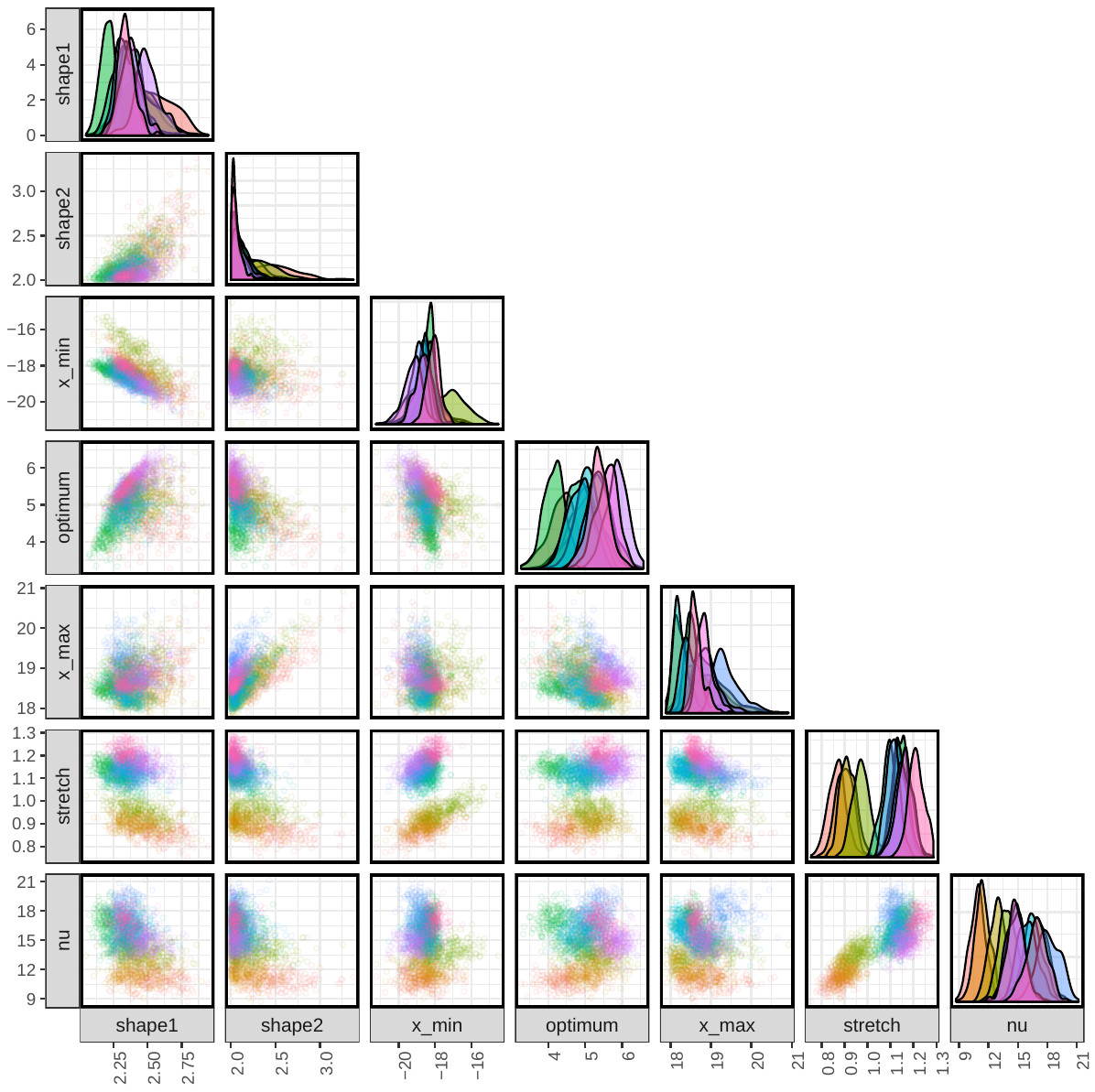

**Figure S3**. Comparison of posterior draws between thermal performance parameters. Parameters are fully described in Tittes et al. (2019), but we briefly describe them here: shape1 and shape2 describe the curve asymmetry (when shape1 is larger than shape2, the curve skews left, and vice versa); xmin and xmax are the temperatures to the left and right (respectively) of the thermal optimum where RGR falls to zero; stretch determines the maximum RGR, or the height of the thermal performance curve; nu describes the variance in RGR. Colors represent different groups as in Fig. S2.

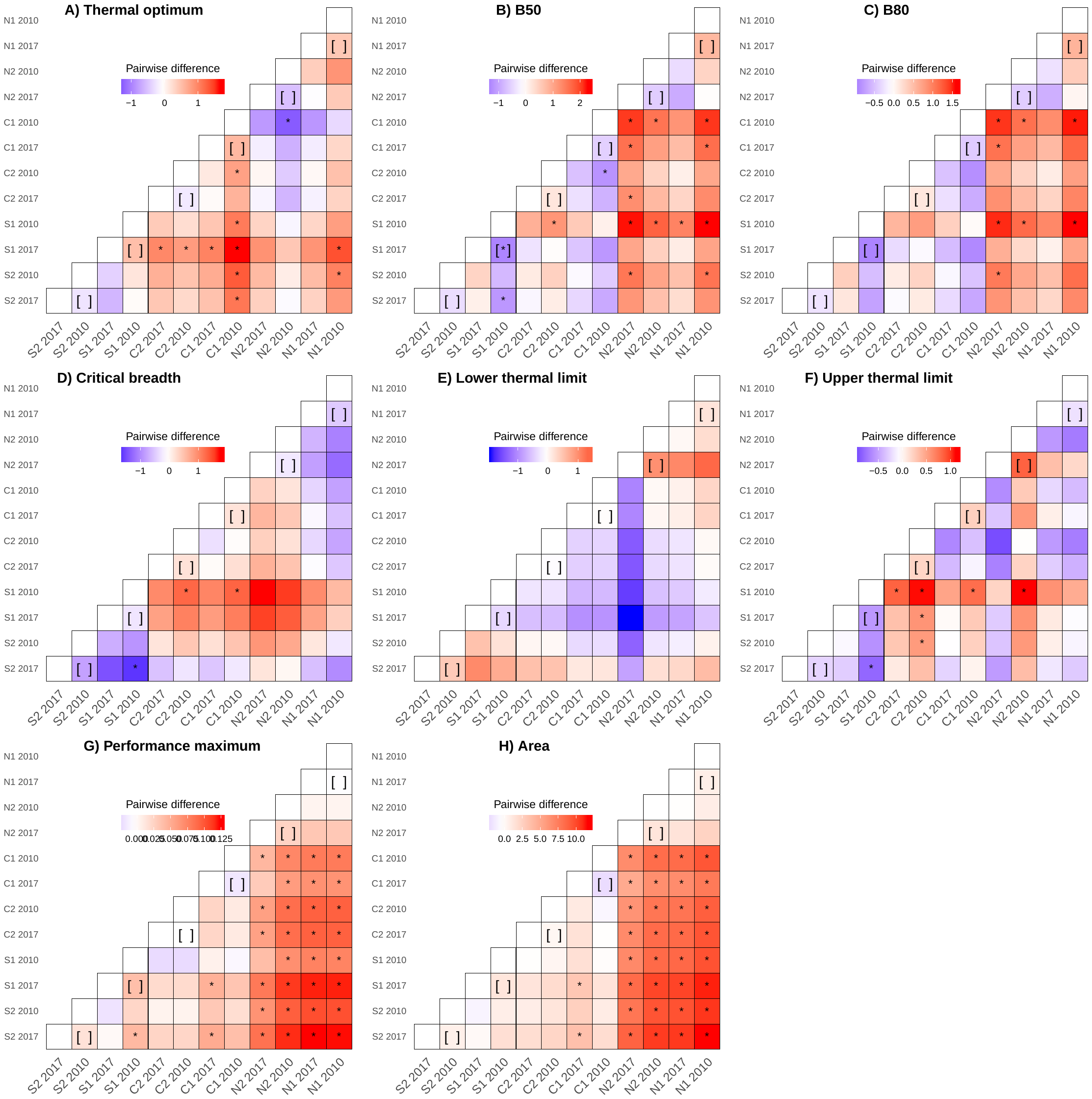

**Figure S4**. Pairwise comparisons of thermal performance parameters across six populations and two cohorts within each population of *Mimulus cardinalis*. Brackets denote comparisons that would indicate evolutionary responses to recent climate change for each population. The color of each box represents the mean pairwise difference between the x group (columns) and the y group (rows) for 20,000 iterations of a Bayesian model. Red indicates cases where the y group has a parameter value that is greater than the x group, and blue indicates cases where the y group has a parameter value that is less than the x group. Asterisks represent pairwise comparisons where the 95% credible interval for differences across all iterations of the model does not overlap zero. Large blocks of asterisks in panels G and H indicate that central and southern populations have higher performance maxima and areas under curves than northern populations. All pairwise differences are in units of °C except performance maximum (leaves leaves^-1^ day^-1^). Parameter descriptions are listed below Table S4.

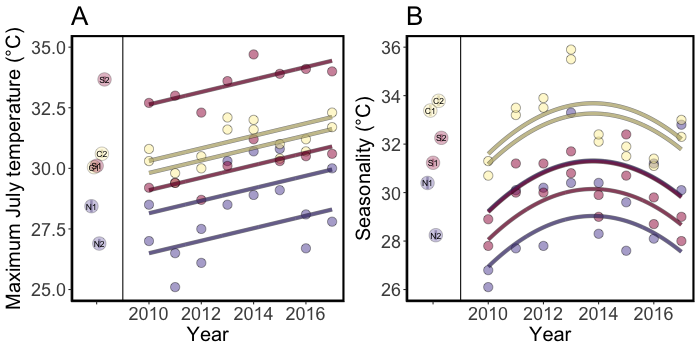

**Figure S5**. Recent climatic trends for two northern-edge (purple), two central (yellow), and two southern-edge (red) populations of *Mimulus cardinalis*. A) Maximum July temperatures increased from 2010 to 2017, whereas B) temperature seasonality increased from 2010 to 2017. Points to the left of regressions represent historical means (± s.e.) for each population. Model specifications and full results are described in Appendix S2. Climate data are described in Table S1.

**Appendix S1.** Supplementary methods for refresher generation and resurrection experiment.

*Refresher generation*

For the refresher generation, we germinated field-collected seeds in pots filled with Fafard 4P Mix (Sun Gro Horticulture, Agawam, MA, USA) and topped with a thin layer of Sunshine Grow Mix (Sun Gro Horticulture, Agawam, MA, USA) in May 2018. Pots were placed into sub-irrigated trays in a greenhouse at North Carolina State University, watered daily, and fertilized weekly with Blossom Booster fertilizer (JR Peters Inc., Allentown, PA, USA) or Cal-Mag fertilizer (Everris, Dublin, OH, USA). Starting at ~4 months after planting, we performed crosses to produce experimental seed families, which were subsequently harvested in September 2018 through January 2019.

*Resurrection experiment*

Beginning in February 2019, we planted seeds into plug trays. Within 24 hours prior to planting, plug trays were filled with Fafard 4P Mix (Sun Gro Horticulture, Agawam, MA, USA), topped with a thin layer of Sunshine Grow Mix (Sun Gro Horticulture, Agawam, MA, USA), and thoroughly misted with water. We planted each family into a separate cell, such that each plug tray contained all ancestral and descendent families of one region (northern, central, or southern) to minimize competition due to size differences among regions. Each set of six trays that eventually went into each growth chamber run contained two representative trays for each region. Due to space limitations, we staggered planting, which occurred through May 2019. Because a previous trial showed that northern populations germinate slower than central and southern populations, we planted the two northern trays in each chamber run one week earlier than the two central and two southern trays to standardize plant size prior to growth chamber runs. Trays were sub-irrigated with a general nutrient solution (containing NPK + micronutrients) once daily. We rotated trays three times per week to reduce positional effects, and emptied water out of trays three times per week to reduce growth of fungus gnat larvae and algae in the soil.

Growth chamber runs occurred from April through July of 2019. After five (for central and southern populations) to six (for northern populations) weeks in the grow room, we put each tray set into growth chambers, with 16 chamber runs total. We randomized the order of temperature regimes and randomly assigned regimes to one of the four growth chambers. During all growth chamber runs, we sub-irrigated trays with reverse osmosis water (as opposed to nutrient solution, which may have confounded any effect of temperature if higher temperatures increased plant water use) twice daily, and randomized trays within each growth chamber twice per week to minimize positional effects.

**Appendix S2.** Analysis of recent trends in maximum July temperatures and temperature seasonality across populations.

The following analyses were performed using simple linear models implemented with the functions *lm* and *anova* from the *stats* package in R (v3.6.1; R Core Team 2015). Climate data are listed in Table S1.

*Maximum July temperature*

To determine if maximum July temperatures varied over the study period from 2010 to 2017, and whether these trends differed between populations, we implemented a model that included maximum July temperatures as the response variable and main effects of year, population, the interaction between year and population, and a second-order effect of year. The simplest model with the lowest AICc score (ΔAICc < 2) included main effects of year and population. This model explained 82% (*R^2^_adj_*) of variation in maximum July temperatures, and indicated that maximum July temperatures increased across the study period (*b*=0.258, *p*<0.001) and varied significantly among populations (*F*_5,41_=39.816, *p*<0.001; Fig. S5A).

*Temperature seasonality*

To determine if temperature seasonality varied across recent years, and whether these trends differed between populations, we implemented a model that included main effects of year, population, the interaction between year and population, and a second-order effect of year (year^2^). The simplest model with the lowest AICc score (ΔAICc < 2) included main effects of year, population, and year^2^. This model explained 61% (*R^2^_adj_*) of variation in temperature seasonality and indicated that temperature seasonality varied significantly among populations (*F*_5,40_= 13.794, *p*<0.001), increased at the beginning of the study period (*b*=1.100, *p*=0.001), and decreased toward the end of the study period (*b*=-0.145, *p*=0.002; Fig. S5B).
